# Lysogenized phages of methanotrophic bacteria show a broad and untapped genetic diversity

**DOI:** 10.1101/2022.05.20.492862

**Authors:** Miranda Stahn, Aurelija M. Grigonyte, Fabini D. Orata, David A. Collins, Liam Rieder, Marina G. Kalyuzhnaya, Andrew Millard, Lisa Y. Stein, Dominic Sauvageau

## Abstract

Methanotrophs are a unique class of bacteria with the ability to metabolize single-carbon compounds such as methane. They play an important role in the global methane cycle and have great potential as industrial platforms for the bioconversion of methane from industrial waste streams into valuable products, such as biofuels and bioplastics. However, many aspects of methanotroph biology have yet to be elucidated, including the prevalence and impact of lysogenized bacteriophages (phages), which can greatly affect both the ecology and the industrial performance of these bacteria.

The present study investigates the presence of putative prophages in three gammaproteobacterial (*Methylobacter marinus* A45, *Methylomicrobium album* BG8, *Methylomonas denitrificans* FJG1) and two alphaproteobacterial (*Methylosinus trichosporium* OB3b, *Methylocystis* sp. Rockwell) methanotrophs using four programs predicting putative phage sequences (PhageBoost, PHASTER, Phigaro, and Island Viewer). Mitomycin C was used to trigger induction of prophages, which was monitored through infection dynamics. Successfully induced phages from *M. marinus* A45 (MirA1, MirA2), *M. album* BG8 (MirB1), and *M. trichosporium* OB3b (MirO1) were isolated and characterized using transmission electron microscopy. Subsequently, bioinformatic analyses (BLAST and phylogenetics) were performed on three induced phages to obtain a profile of their respective genetic makeup. Their broad diversity and differences from previously known phages, based on whole genome and structural gene sequences, suggest they each represent a new phage family, genus and species: “*Britesideviridae Inducovirus miraone*”, “*Patronusviridae Enigmavirus miratwo*”, and “*Kainiviridae Tripudiumvirus miroone*” represented by isolates MirA1, MirA2, and MirO1, respectively.

## Introduction

Methanotrophic bacteria can utilize single-carbon compounds such as methane as their sole source of carbon and energy. They play an important role in the global methane cycle and show promise as industrial platforms for the conversion of methane from impure sources, such as industrial waste streams, into a wide array of valuable products ranging from isoprenoids to polyhydroxyalkanoates and ectoines, among others (Cantera et al., 2018; Henard et al., 2017; Jiang et al., 2010; Khmelenina et al., 1999; Lee et al., 2016; Strong et al., 2015). Despite first being characterized in the early 1900s (Kaserer, 1905; Söhngen, 1906), much about methanotroph biology has yet to be elucidated, including the prevalence and impact of bacteriophages (phages) in these microorganisms.

Considering the ubiquity of phages in nature, phages of methanotrophs have been assumed to exist and have been included in ecological models of microbial communities (Emerson et al., 2018) and identified in metagenomic studies (Gambelli et al., 2016). Recently, transcriptomics data from Lake Rotsee (Switzerland) was used to suggest some phages were found to encode the *pmoC* gene, a subunit of methane monooxygenase, which is the prevalent methane oxidation catalyst in nature (Chen et al., 2020). The research suggested that *pmoC*-containing phages infecting methanotrophs could directly impact methane oxidation rates and thus methane emissions. Another study, focusing on in situ ^13^C-enriched metabolomics of soil bacterial communities, demonstrated the potential role of phage predation on the release of methane-derived compounds in soils (Lee et al., 2021). But until now, only two studies, both from the early 1980’s, have reported the isolation and partial characterization of (strictly lytic) phages of methanotrophs (Tyutikov et al., 1980; Tyutikov et al., 1983).

Since temperate phages are generally considered more abundant than strictly lytic phages (Angly et al., 2006), lysogenized phages are of particular interest. This has two significant implications. For one, despite being a metabolic burden on the cell (Ramisetty & Sudhakari, 2019), the presence of a prophage in the host genome can confer an evolutionary advantage to this host – e.g., encoding genes for multi-drug resistant pumps, outer membrane proteases, or toxic membrane polypeptides (Ramisetty & Sudhakari, 2019; Wang & Wood, 2016). Secondly, the induction of a prophage can lead to rapid changes in microbial communities in a given environment, or to premature cell lysis and the subsequent loss of an entire production batch in industrial applications – a problem often encountered in the dairy industry (De Paepe et al., 2016; Kilic et al., 1996).

The lack of information on temperate phages of methanotrophs may be a side-effect of the limited amount of information available for this group of bacteria. But the increasing number of sequenced methanotroph genomes and the advancement of bioinformatics tools used for prophage prediction – e.g., PhageBoost (Sirén et al., 2021), PHASTER (Arndt et al., 2016; Zhou et al., 2011), and Phigaro (Starikova et al., 2020) – provide new platforms for the investigation of these phages.

In the present study, temperate phage induction was attempted in five methanotrophic bacteria: *Methylobacter marinus* A45, *Methylomicrobium album* BG8, *Methylomonas denitrificans* FJG1, *Methylosinus trichosporium* OB3b, and *Methylocystis* sp. Rockwell. Successfully induced phages were characterized through microscopy and genetic analysis. Comparisons were made with putative phage sequences predicted using bioinformatic tools. Phylogenetic and alignment analyses of selected phage genes were also used to reveal the great genetic diversity of the induced phages. This study provides information on the prevalence of prophages in methanotrophic bacteria, describes four temperate phages of methanotrophs, and demonstrates the broad genetic diversity in these phages.

## Materials and Methods

### Microorganisms and Cultures

Three gammaproteobacterial methanotrophs (*Methylobacter marinus* A45, *Methylomicrobium album* BG8 (ATCC 33003), and *Methylomonas denitrificans* FJG1) and two alphaproteobacterial methanotrophs (*Methylosinus trichosporium* OB3b and *Methylocystis* sp. Rockwell (ATCC 49242) were used in this study. Cultures were maintained by inoculating 100 ml of ammonium mineral salt (AMS) or nitrate mineral salt (NMS) media (Whittenbury, Phillips, & Wilkinson, 1970) with 1 ml of methanotroph culture in early stationary phase of growth into a 250-ml Wheaton bottle sealed with a screw-top cap inlaid with a butyl rubber septum. 50 ml of headspace gas was removed using a gas-tight syringe and replenished with 60 ml of methane (Praxair, Edmonton, Canada), pressurizing the bottle to facilitate methane diffusion into the medium. To facilitate the growth of the oceanic strain *M. marinus* A45, 100 ml of 1X NMS was supplemented with 6.65 ml 1% NaCl, 2 ml phosphate buffer, and 0.5 ml 1 M sodium bicarbonate solution. Cultures were incubated at 30°C with shaking at 150 rpm. Cell density was assessed by measuring optical density at 540 nm (OD_540_) using a UV-Vis spectrophotometer (Biochrom Ultropsec 50).

### Prophage Induction

Bacterial cultures (in 50 ml medium as described above) were grown to mid-log phase (OD_540_ ∼0.15-0.30), at which point MitC (Sigma-Aldrich, USA) was added to cultures to a final concentration of 1 or 2 μg/ml. Cultures were left to grow in the presence of MitC for 1 h, with the Wheaton bottles wrapped in tinfoil to protect from light. Following exposure, 30 ml of culture was centrifuged for 20 min at 20,000 × g and 20°C (Sorvall Evolution RC, Kendro Laboratory Products, USA) to remove MitC from the medium. Cell pellets were resuspended in 30 ml of fresh medium, the methane was replenished to the headspace, and cultures were left to grow for ∼2 days. OD_540_ was measured periodically to monitor culture growth and potential prophage induction (signified by a reduction in OD_540_). Other signs of prophage induction included clumping, color change of the culture, and precipitation or debris in the medium, which were used concomitantly with OD to determine successful induction.

### Phage Isolation and Characterization

Cultures showing signs of induction were passed through a 0.2-μm pore size PTFE syringe filter to remove whole cells and debris from lysis. Filtered lysates were stored at 4°C. To prepare samples for transmission electron microscopy (TEM), 4 ml of cell lysate were centrifuged at 20,000 × g at 4°C for 1 h. The supernatant was discarded, and the pellets were resuspended in 1 ml of lambda diluent (Roskams and Rodgers, 2002) and left to settle overnight at 4°C. Pellets were concentrated by centrifuging under the above conditions, removing the supernatant and resuspending in 100 μl of lambda diluent. 10 μl of the sample were then placed on a copper-coated grid/Formvar film, left to sit for 4 min, and then stained with uranyl acetate for 1 min. Using a Morgagni 268 TEM (FEI, Hillsboro, Oregon, USA), samples were visualized at 110,000× magnification, using a beam intensity of 80 kV. Phage capsid and tail sizes were calculated by measuring a minimum of five samples in ImageJ.

Phage DNA was recovered from filtered lysate. 50 ml of lysate was aliquoted into an Eppendorf tube and spun at 20,000 × g at 4°C for 2 h. The supernatant was then discarded and samples were resuspended in 300 μl of SM (NaCl (100 mM) MgSO4·7H_2_O (8mM) Tris-Cl (50 mM)) buffer (Puapermpoonsiri et al., 2010) and allowed to settle overnight. Samples were then centrifuged again for 20,000 × g at 4°C for 1 h, the supernatant was replaced with 10 μl phage buffer, and the samples were recombined as a single volume. Samples were centrifuged once more at 20,000 × g for 1 h and resuspended into 200 μl of buffer for extraction using a DNA isolation kit (Norgen, Canada). Each 20 μl of DNA sample isolated per 50 ml lysate were pooled, creating a master stock for sequencing using a MinION nanopore sequencer (Oxford Nanopore Technologies, Oxford, UK).

### Library preparation for MinION sequencing

A total of ∼1 μg purified DNA from each induced prophage sample was used to prepare sequencing libraries using a Ligation Sequencing Kit 1D (SQK-LSK109) and Native Barcoding Kit 1D (EXP-NBD104) following manufacturer’s instructions. Pooled libraries were sequenced on a MinION Flow Cell (R9.51) with data processed on a MIN-IT (Oxford Nanopore). Base calling was done with guppy v2.1 using “--flowcell FLO-MIN106 --kit SQK-LSK109” settings. To identify induced prophage regions, reads were mapped against the reference genome with Minimap2 v.2.14 with the following settings ‘-a -x ont-map -t’. Coverage of prophage regions was determined using samtools v1.6 to calculate coverage over a 500 bp window to compare against the background level of bacterial DNA.

### Prophage Prediction

Methanotroph genome sequences used in this study were acquired from GenBank (accession nos. ARVS00000000, CM001475, CP014476, ADVE00000000, AEVM00000000).

The sequences were analyzed through the prophage predictor software programs PHAge Search Tool – Enhanced Release (PHASTER) (Arndt et al., 2016; Zhou et al., 2011), Phigaro v2.3.0 (Starikova et al., 2020), PhageBoost v0.1.7 (Sirén et al., 2021) and the genomic island identifier IslandViewer 4 (Bertelli et al., 2017), all used with default parameters.

### Genetic Analysis of Induced Phage Sequences

Phage genome sequences obtained from MinION sequencing were annotated with Prokka v1.14.6 (default settings) using a custom database of all phage genomes extracted from GenBank (Seemann, 2014; Cook et al., 2021). Further analysis and annotation were carried out using the prokaryotic virus orthologous groups (pVOGs) database to identify any proteins within pVOGs using hmmscan (default settings) (Eddy, 2011; Grazziotin et al., 2017). Raw sequence data and assembled genomes were deposited in the European Nucleotide Archive (ENA) under the project accession number PRJEB46923.

Basic local alignment analysis was performed using the NCBI Basic Local Alignment Search Tool (BLAST) for whole phage genomes (based on nucleotide sequences; nBLAST) and for selected phage gene products (based on protein sequences; pBLAST) of induced phages against both the NCBI complete microbial and *Caudovirales* (taxid:28883) databases. Sequence alignment was performed using the MAFFT v7.271 (default settings) multiple sequence alignment program (Katoh & Toh, 2008). Both the percent coverage and percent identity were reported.

Phylogenetic analyses were carried out for selected phage genes (major capsid and tail-associated proteins). Homologues for each were obtained from the NCBI GenBank database for phages from known bacterial hosts and the top ten hits (by BLAST score) from the IMG/VR database (Roux et al., 2021) for uncultivated viruses. The amino acid sequences were first aligned with MUSCLE v3.8.425 (Edgar, 2004). The alignments were then used to reconstruct maximum-likelihood phylogenetic trees using RAxML v8.2.12 (Stamatakis, 2014). The WAG (Whelan and Goldman) amino acid substitution model and gamma model of rate heterogeneity were used. Robustness of branching was estimated with 1,000 bootstrap replicates. Tree patristic distances were determined using Geneious v11.1.5 (Kearse et al., 2012). The trees were visualized using the Interactive Tree of Life (iTOL) v6.3 (Letunic & Bork, 2021). The phylogenetic trees for the entire induced phage genomes were carried out using VipTree software version 1.9 (default parameters for dsDNA prokaryotic viruses) (Nishimura et al., 2017).

The induced prophage sequences were also analyzed for virulence factors and genes associated with antibiotic resistance using VirulenceFinder 2.0 (Tetzschner et al., 2020) and ResFinder 4.0 (Bortolaia et al., 2020), using the default parameters.

## Results

### Induction of prophages

The induction of prophages was monitored through dynamic OD_540_ measurements of batch cultures following addition of MitC to 1 and 2 μg/ml, with a reduction in OD_540_ indicative of potential cell lysis, and hence prophage activation. The area under the curve of OD_540_ (AUC) was then used to assess the significance of lysis between each experimental condition against the control (host culture growing in the absence of MitC). A lower AUC value is indicative of reduced growth and, consequently, reduced viability. Figure 1 shows results for *M. marinus* A45, *M. album* BG8, *and M. trichosporium* OB3b. Despite showing “symptoms” of induction (i.e., precipitation, cell aggregates, and lower OD compared to controls), TEM analysis could not confirm the presence of induced phage in cultures of *Methylocystis* sp. Rockwell, and *M. denitrificans* FJG1; the data for these strains can be found in the Supplementary Materials (Supplementary Figure S1).

**Figure 1.**
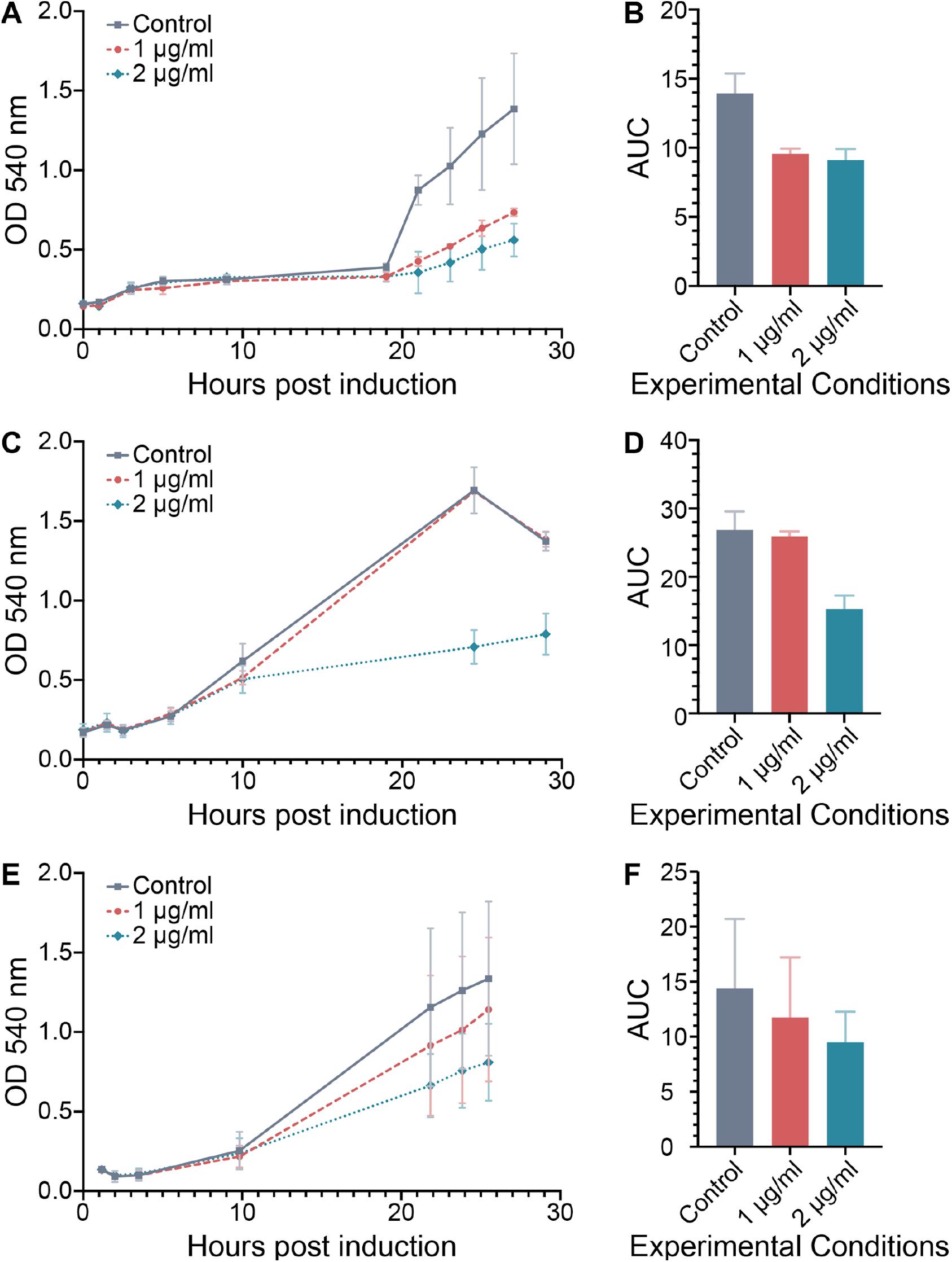
Induction of potential prophages by addition of mitomycin C to cultures of different methanotrophs. Optical density (OD_540_) and the area under the curve of OD_540_ (AUC) are shown for *M. marinus* A45 (A, B), *M. album* BG8 (C, D), and *M. trichosporium* OB3b (E, F) after the addition of 1 μg/ml and 2 μg/ml of mitomycin C and control experiments. Error bars denote the standard deviation based on n = 3.

In all but one case, bacterial growth was negatively affected by 1 μg/ml MitC (Figures 1A, 1E and Supplementary Figures S1A, S1C). *M. album* BG8 was the only exception, whereby growth was only impeded at 2 μg/ml MitC (Figure 1C). AUC and two-way ANOVA analysis were used to validate the statistical significance of these results. *M. marinus* A45, *M. denitrificans* FJG1 and *Methylocystis* sp. Rockwell showed significant differences in cell viability in the presence of 1 μg/ml MitC compared to the control, exhibiting p-values <0.005 (Figure 1B and Supplementary Figures S1B and S1D). On the other hand, all cultures showed a significant difference between the 2 μg/ml MitC condition and the controls (Figures 1B, 1D, and 1F, and Supplementary Figures S1B and S1D) – with p-values <0.0001 for all strains, except *M. trichosporium* OB3b which had a p-value of 0.0370. This suggests a reduction in culture viability likely indicative of successful prophage inductions.

Since none of the induction experiments demonstrated the characteristic “crash” in OD_540_ resulting from population-wide lysis typical of virulent infections, additional measures were used to characterize potential induction. Inductions were considered successful if any of the following was observed: 1) a greater than two-fold difference between the final OD_540_ of the control and the experimental subset – final OD_540_ of the control was based on the time at which the culture reached stationary phase; and/or 2) signs of disrupted or unhealthy cultures, such as precipitation of cell debris/dead cells.

### Characterization of induced phages

Experimental samples presenting at least two conditions linked to potential prophage induction were analyzed under TEM. Phages were detected in samples from *M. marinus* A45, *M. album* BG8, and *M. trichosporium* OB3b and designated as isolates MirA1, MirB1, and MirO1, respectively (Figure 2). The main physical characteristics of the recovered phages can be found in Table 1. No phages were observed in samples from cultures of *Methylocystis* sp. Rockwell or *M. denitrificans* FJG1.

**Figure 2.**
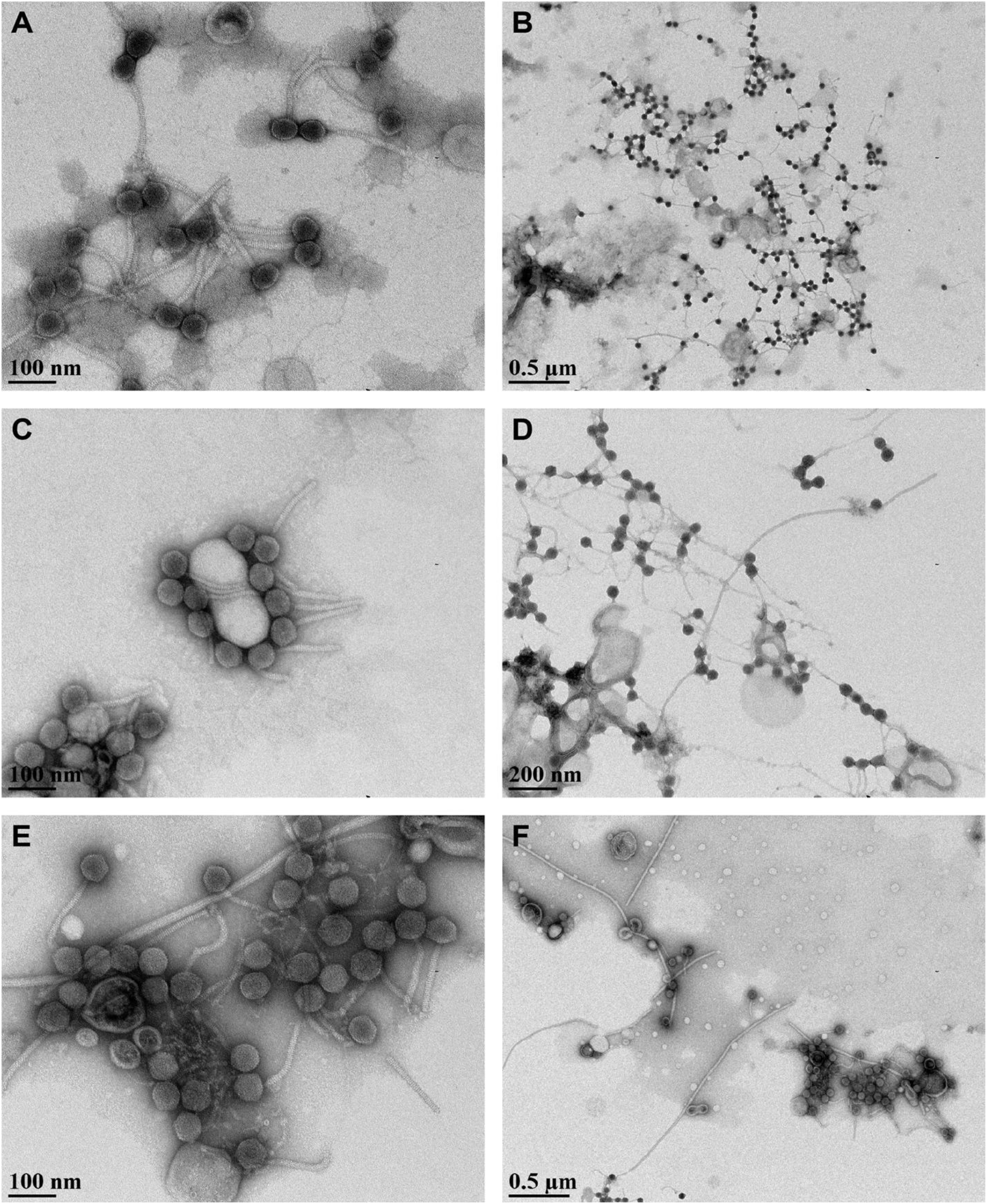
Transmission electron microscopy visualization of induced cultures of *M. marinus* A45 (A, B), *M. album* BG8 (C, D), and *M. trichosporium* OB3b (E, F) using uracil acetate staining.

**Table 1.**
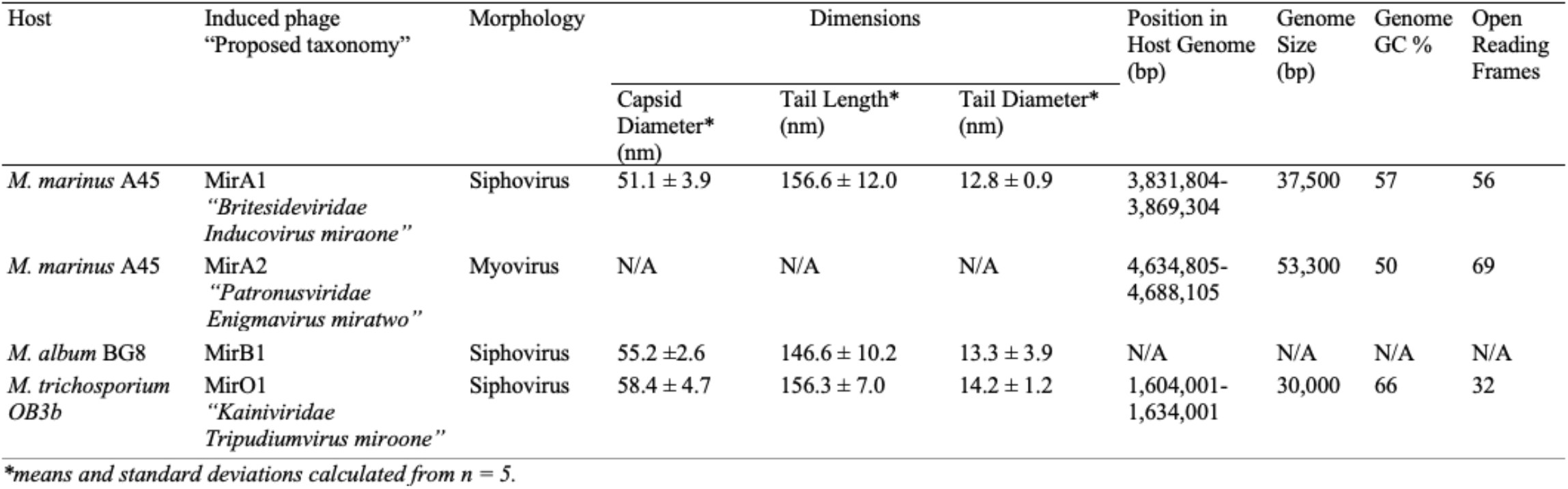
Properties of phages induced from *M. marinus* A45, *M. album* BG8, and *M. trichosporium* OB3b.

All three induced phages displayed Siphovirus morphology – symmetrical capsid, long flexible tails. They also had similar dimensions, with capsid diameters ranging between ∼51-58 nm, and their long helical tails ranging between ∼146-157 nm in length and ∼13-14 nm in diameter (Table 1).

### Prediction of prophage regions

Four different software packages – PHASTER, Phigaro, PhageBoost and IslandViewer – were used to detect and cross-analyze prophage regions from the five methanotrophs of interest – *M. marinus* A45 (Figure 3A), *M. album* BG8 (Figure 3B), *M. trichosporium* OB3b (Figure 3C), *M. denitrificans* FJG1 (Supplementary Figure S2A), and *Methylocystis* sp. Rockwell (Supplementary Figure S2B). In the cases where the methanotroph genome was comprised of two or more contigs (*M. marinus* A45 and *Methlyocystis* sp. Rockwell), each genome segment was run through the algorithms individually. Only contigs that showed putative prophage regions are shown in the results.

**Figure 3.**
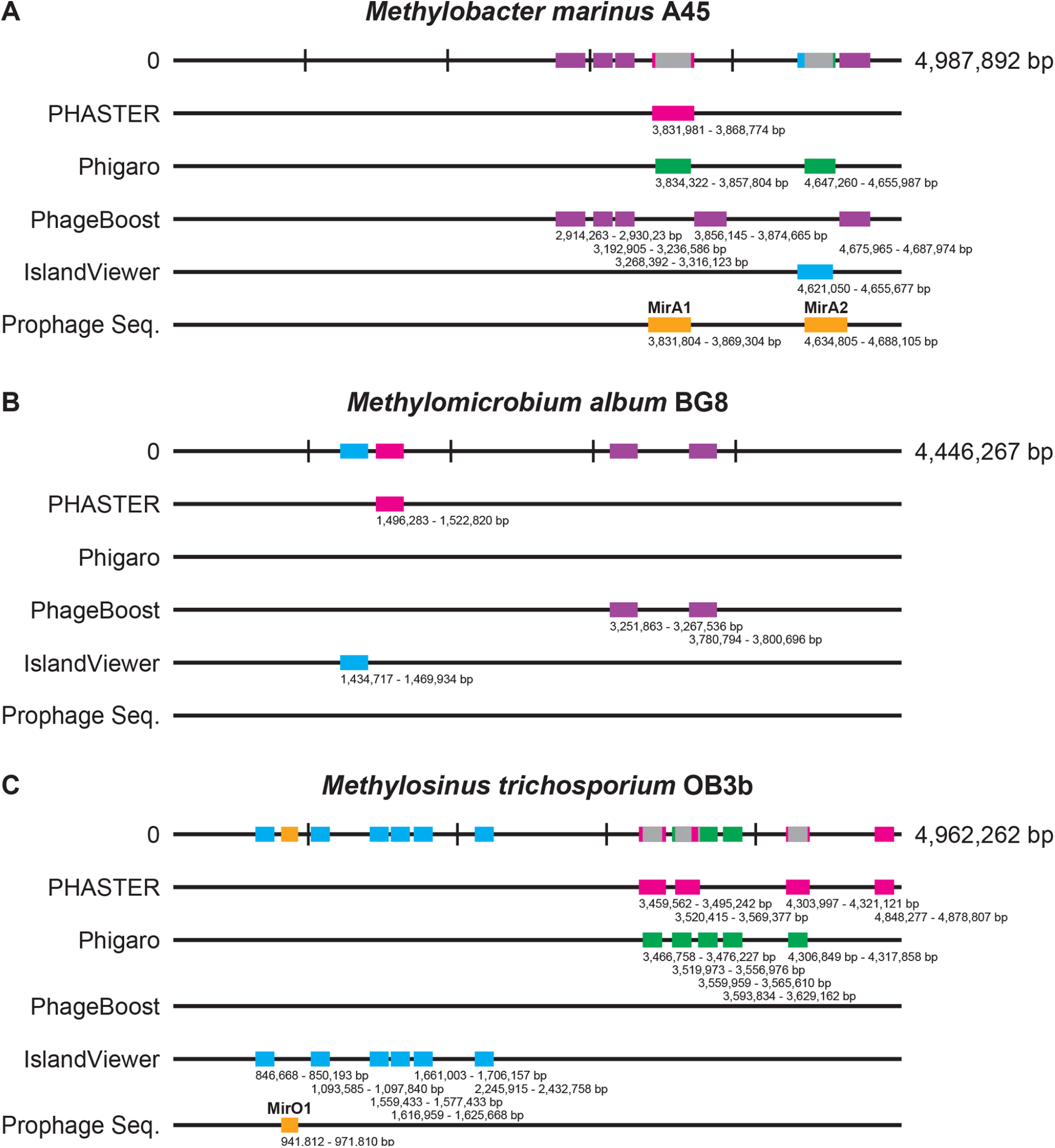
Computational predictions of prophage regions using PHASTER, Phigaro, PhageBoost, and IslandViewer in the gammaproteobacterial methanotrophs *M. marinus* A45 (A), *M. album* BG8 (B), and *M. trichosporium* OB3b (C). Reference genome sequences are reported at the top of the figures, followed by visualizations of putative prophage regions identified by PHASTER (magenta), Phigaro (green), PhageBoost (violet), and IslandViewer (blue). Overlapping putative regions are illustrated in grey in the host genome when two predicted regions overlap. Sequences of isolated induced phages are shown at the bottom of the figures (yellow). The location of each region of interest is indicated by the base pair numbers corresponding to their position in the host genome.

All four algorithms detected prophage regions in *M. marinus* A45 (Figure 3A). One region, starting in the 3,830,000 bp area, was identified by both PHASTER and Phigaro, albeit with slightly different start and end points. This region also corresponded to the genetic sequence of induced prophage MirA1 (Table 1). Interestingly, the ends of the induced phage sequence were very close to those predicted by Phigaro. Another region was co-identified by Phigaro and IslandViewer in low end of the 4,600,000 bp area. This region also corresponded to the genetic sequence of an induced prophage, MirA2 (Table 1).

The same exercise performed on the genome sequence of *M. album* BG8 identified four different putative regions (one from PHASTER, two from PhageBoost, and one form IslandViewer) with no overlap between algorithms (Figure 3B). Phigaro did not detect putative prophage regions in the genome of *M. album* BG8.

In *M. trichosporium* OB3b, a large number of putative regions were identified by PHASTER, Phigaro and IslandViewer, but none by PhageBoost (Figure 3C). Two overlapping regions were found by PHASTER and Phigaro in the 3,460,000 bp and 3,520,000 bp areas. Phage DNA isolated from induction experiments showed a 30,000 bp genome region corresponding to the host sequence at 941,812 – 971,810 bp (Table 1). This region did not overlap with any of the regions predicted by the four software programs.

In *M. denitrificans* FJG1, only Phigaro (one) and PhageBoost (four) detected putative prophage regions (Supplementary Figure S2A). No induced phages were observed in the induction experiments. The genome of *Methylocystis* sp. Rockwell was divided into six different contigs (each corresponding to an individual GenBank file). The algorithms only detected putative prophage regions contig MAH.6 (Supplementary Figure S2B). A large number of putative prophages were detected, with three overlaps (between Phigaro and IslandViewer in the 1,680,000 bp area; between PHASTER and IslandViewer in the 2,660,000 bp area; and between PHASTER and Phigaro in the 3,420,000 bp area). Here again, no induced phages were observed in the induction experiments.

### Genetic diversity of induced phages

To better understand the genetic make-up and genome architecture of the induced phages, the genomes of phages MirA1, MirA2 and MirO1 were sequenced and annotated. Their respective genetic characteristics are found in Table 1. Complete genome sizes varied from 30 to 53.3 kb and comprised between 0.57 to 1.06% of the host chromosome. The percentage of hypothetical genes was 62.5%, 66.6%, and 50% for phages MirA1, MirA2, and MirO1, respectively.

Figure 4 shows the genome composition with annotation (including encoded protein direction) for these three phages. In the case of MirA1, only one out of fifty-six encoded proteins, putative cI repressor, was located on the 5’ to 3’ strand (Figure 4A). In contrast, MirA2 had a mixture of bidirectionally encoded proteins where 26/69 genes were found on the 5’ to 3’ strand (Figure 4B), while MirO1 had 31/32 encoded proteins on the 5’ to 3’ strand (Figure 4C). The genes for the main phage structural components (e.g., tail and capsid proteins) were identified for all three phages (Figure 4). Virion proteins were only identified in the MirA1 genome, whereas holin was found only in the genome of MirA2. Interestingly, proteins encoding for transcriptional regulators were identified in both phages induced from *M. marinus* A45. LacI-like regulatory protein and cI repressor protein were found in MirA1, whereas a LuxR family and an unnamed transcriptional regulator protein were found in MirA2. None of the induced phages were found to harbor genes encoding for antibiotic resistance or allergens/toxins when analyzed by VirulenceFinder and ResFinder. None of the prophages were collocated with *pmo*-gene clusters and they did not carry *pmo*C homologues or any proteins with a function associated to the central pathway for methane oxidation and/or assimilation.

**Figure 4.**
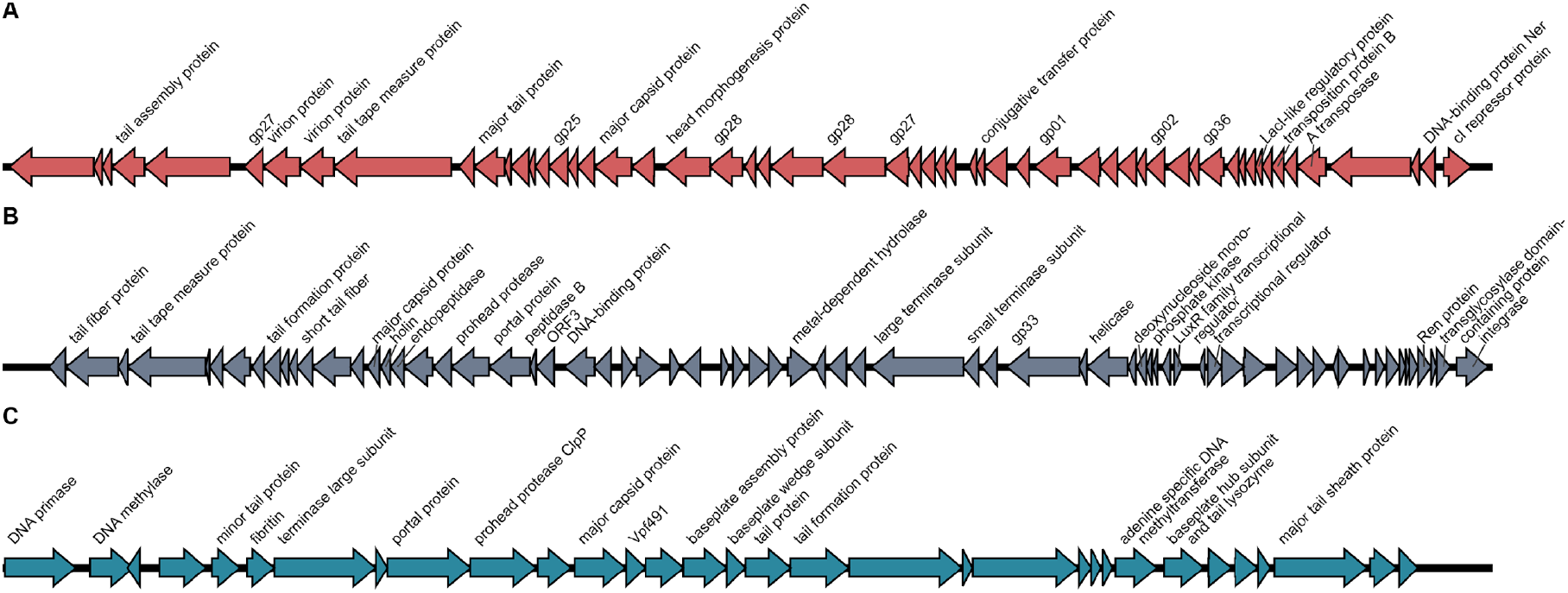
Genome annotations of induced prophages MirA1 (A), MirA2 (B), and MirO1 (C).

As little is known about phages of methanotrophic bacteria, BLAST analysis was performed for the induced phage genomes (nBLAST) and selected relevant annotated phage proteins (pBLAST) against the entire NCBI microbial and *Caudovirales* (taxid:28883) databases. This analysis aimed to provide information on homology with known phages and phage proteins, and shed light on the diversity of phages of methanotrophs and the origin of their proteins.

The top five hits for maximum coverage of the genomes and selected genes of interest are summarized for induced phages MirA1, MirA2, and MirO1 in Figures 5-7 (whole genome, major capsid protein, and tail protein genes) and Supplementary Figures S3-S5 (other genes of interest). For each sequence analyzed (genome or gene), the maximum coverage and percent identity were reported for each of the top five hits. Overall, 100% coverage hits had between 99.79% and 21.07% identity across NCBI and *Caudovirales* pBLAST outputs. In most instances of gene product analysis, the output of pBLAST against the entire NCBI database was dominated by hypothetical proteins found in bacterial host strains rather than phage-specific proteins (i.e., reported from a known phage genome).

**Figure 5.**
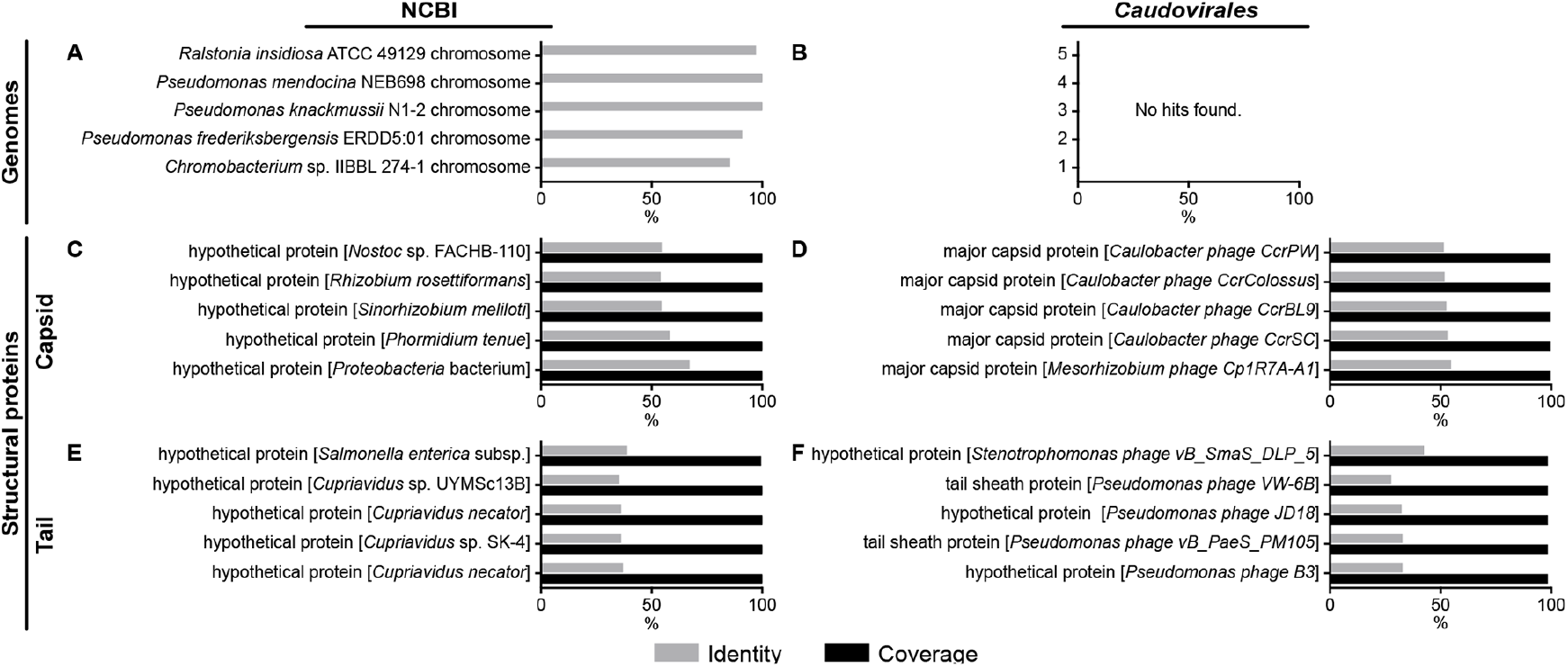
BLAST analysis of induced prophage MirA1 genome (A, B), main capsid protein (C, D) and main tail protein (E, F). Sequences were analyzed against the entire NCBI (A, C, E) or *Caudovirales*-only (B, D, F) databases. Percent identity (gray) and percent coverage (black) are reported. The top five hits based on percent identity were ordered based on percent coverage. Only percent identity is shown for whole genome analyses (A, B) since the coverage was < 3%.

**Figure 6.**
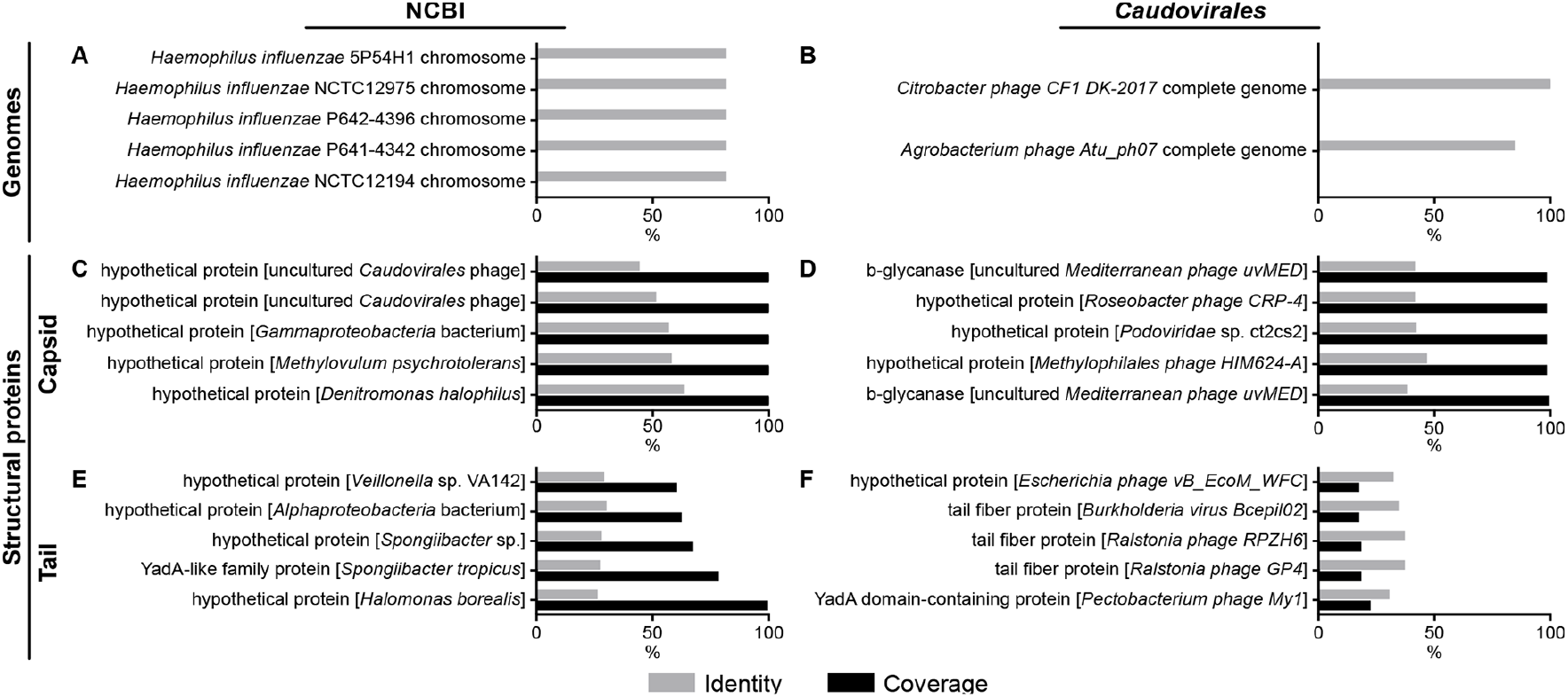
BLAST analysis of induced prophage MirA2 genome (A, B), main capsid protein (C, D), and tail fiber protein (E, F). Sequences were analyzed against the entire NCBI (A, C, E) or *Caudovirales*-only (B, D, F) databases. Percent identity (gray) and percent coverage (black) are reported. The top five hits based on percent identity were ordered based on percent coverage. Only percent identity is shown for whole genome analyses (A, B) since the coverage was < 3%.

**Figure 7.**
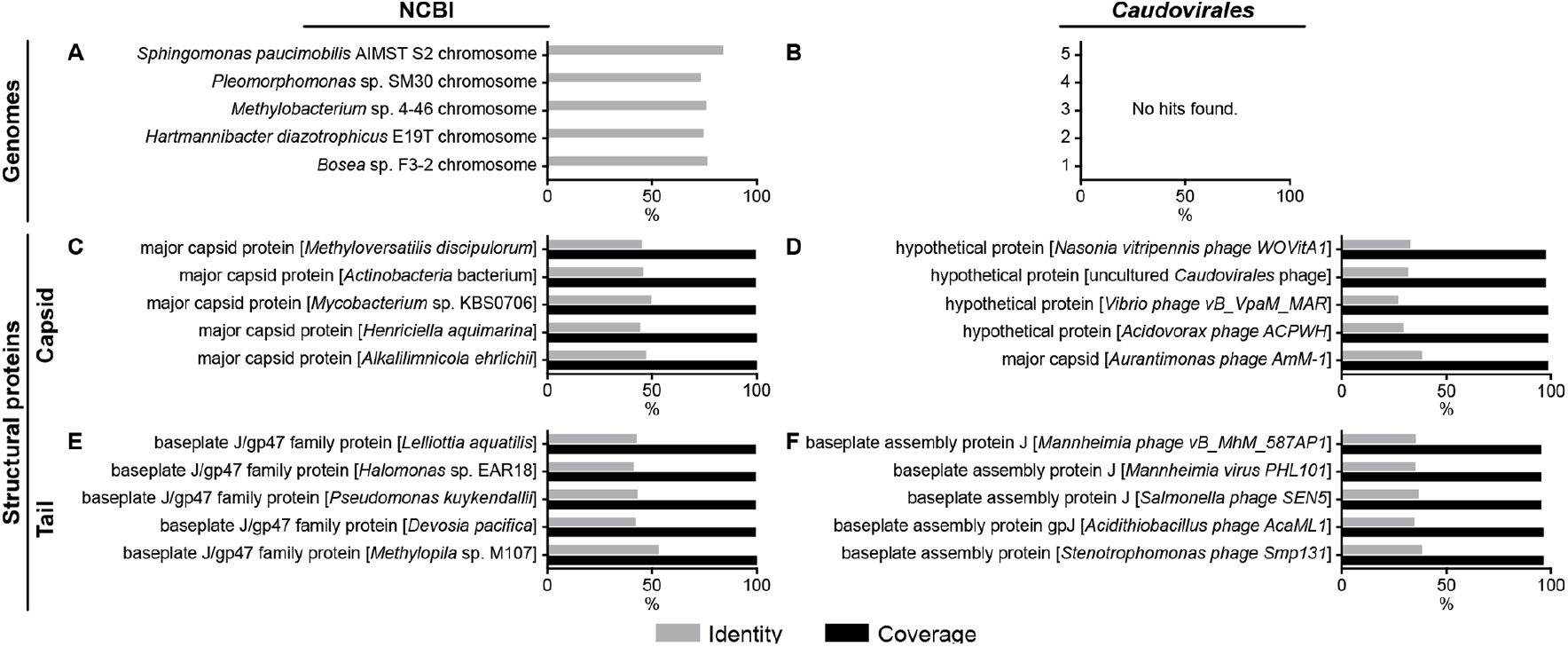
BLAST analysis of induced prophage MirO1 genome (A, B), main capsid protein (C, D), and tail fiber protein (E, F). DNA sequences were analyzed against the entire NCBI (A, C, E) or *Caudovirales*-only (B, D, F) databases. Percent identity (gray) and percent coverage (black) are reported. The top five hits based on percent identity were ordered based on percent coverage. Only percent identity is shown for whole genome analyses (A, B) since the coverage was < 3%.

nBLAST analysis of the phage MirA1 genome (Figure 5A) against the NCBI database identified insignificant coverage (less than 1%) for all top five hits, including sequences from multiple bacterial strains. No significant similarities were found when screening the *Caudovirales*-only database (Figure 5B). This is important as it suggests this phage is representative of a new species, new genera and new family of phage. The current agreed upon cutoffs of 95% ANI for phage species and 70% identity over 100% of the genome (Turner et al. 2021) are clearly met.

The gene products of interest from phage MirA1 for which pBLAST analyses were performed are summarized in Figure 5C-F and Supplementary Figure S3. Again, these analyses generally showed low percent identity (below 55% in all but one case) with sequences from a broad range of bacterial strains (from the NCBI database; Figures 5C, 5E, S3A) and phages of a broad range of hosts (from the *Caudovirales* database; Figures 5D, 5F, S3B).

All nBLAST analysis of the phage MirA2 genome resulted in hits of less than 1% maximum coverage (five hits to *Haemophilus influenzae* genomes from NCBI microbial database, and two hits of *Citrobacter* phage CF1 DK-2017 and *Agrobacterium* phage Atu_ph07 from the *Caudovirales* database; Figure 6A-B). Here again, phage MirA2 meets the requirements for new phage species, genera and family.

The gene products of interest from phage MirA2 for which pBLAST analyses were performed are summarized in Figure 6 and Supplementary Figure S4. Although all five top hits from the NCBI database for the capsid protein are hypothetical proteins, it is interesting to see that one of these (with 55% identity) is from the gammaproteobacterial methanotroph *Methylovulum psychrotolerans* (Figure 6C). When comparing with the *Caudovirales* database, all top hits to the capsid protein have identities below 50%. The scores for the MirA2 tail fiber protein are even lower with coverage between 60-100% for the NCBI database hits and below 25% for the *Caudovirales* database hits. Although three of the top hits from the latter are tail fiber proteins, the coverage reported is low and the identities are all below 45%. Again, a broad variety of phage host is found in the top hits.

Much like the other two phages described above, nBLAST analysis of the phage MirO1 genome against the full NCBI microbial database showed 2% and 1% maximum coverage (Figure 7A). There were no significant similarities when screening the MirO1 genome against the *Caudovirales* database (Figure 7B). Like the two previous phages, MirO1 then also meets the requirements for new phage species, genera and family.

The gene products of interest from phage MirO1 for which pBLAST analyses were performed are summarized in Figure 7C-F and Supplementary Figure S5. Although the coverages were all high (at or near 100%), none of the hits displayed a % identity above 50%. Of interest is also the fact that the hits for the tail fiber protein from the *Caudovirales* database were baseplate proteins.

Further genomic analysis was performed. MirA1, MirA2, and MirO1 genomes were screened (nBLAST) against the IMG/VR database with no significant hits identified. Furthermore, the genomes were used to construct phylogenetic trees (Supplementary Figures 6-8) that revealed little to no clustering and long branching with their closest relatives.

Phylogenetic trees were reconstructed for the genes coding for the major capsid protein and a tail-associated protein of each induced phage: MirA1, MirA2, and MirO1 (Figures 8-10).

**Figure 8.**
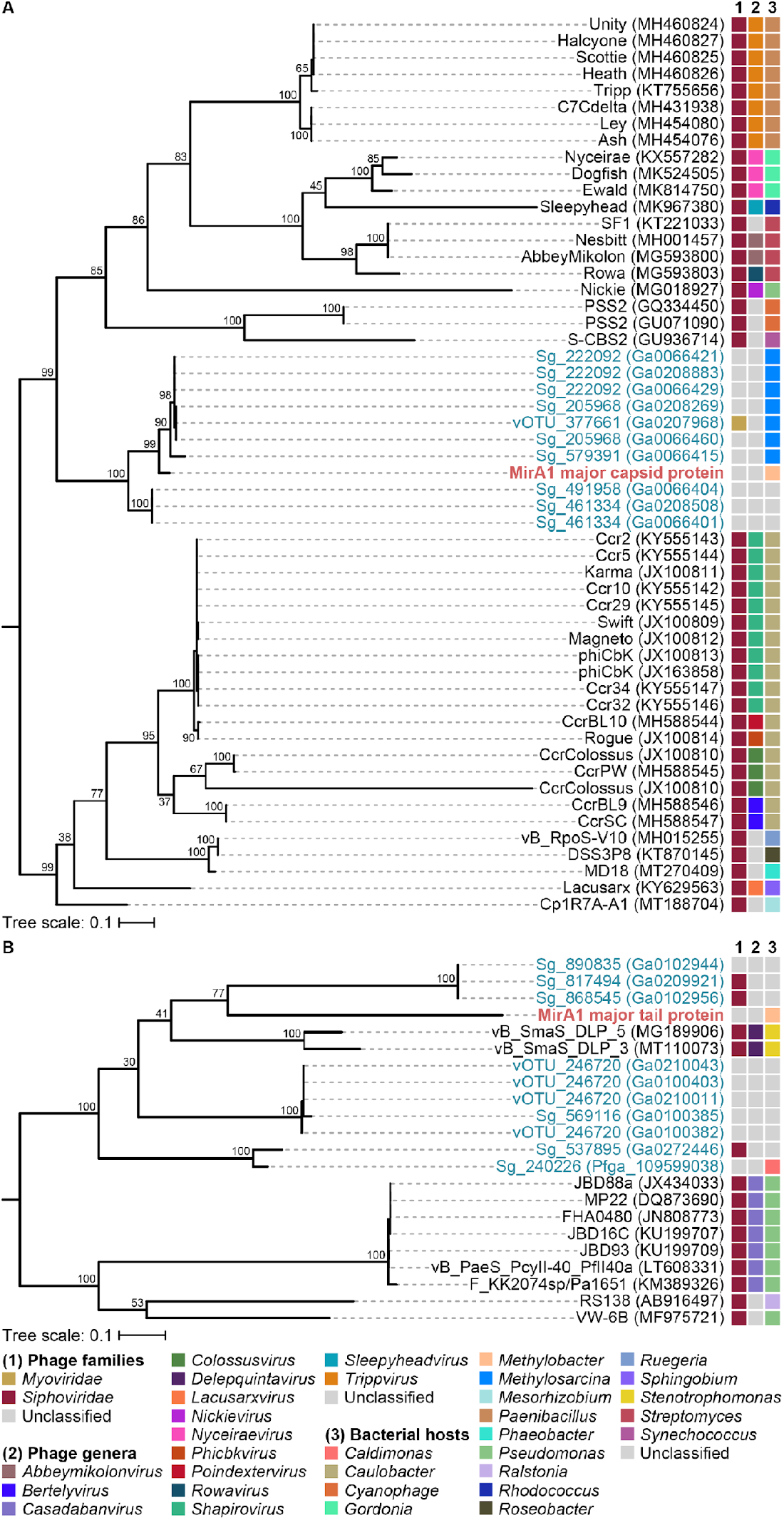
Phylogenetic analysis of induced phage MirA1 major capsid (A) and tail-associated (B) proteins. Midpoint-rooted, maximum-likelihood trees were reconstructed from amino acid sequences (254 and 234 amino acid positions, respectively). Bootstrap support is indicated on the nodes. The scale bar represents amino acid substitutions per site. Tree tip colors indicate genes from the induced phage in this study (red), from NCBI hits (black), or from uncultivated sources (blue). Additional annotations include known (for NCBI hits) or predicted (for IMG/VR hits) 1) phage family designations, 2) phage genus designations, and 3) bacterial hosts.

**Figure 9.**
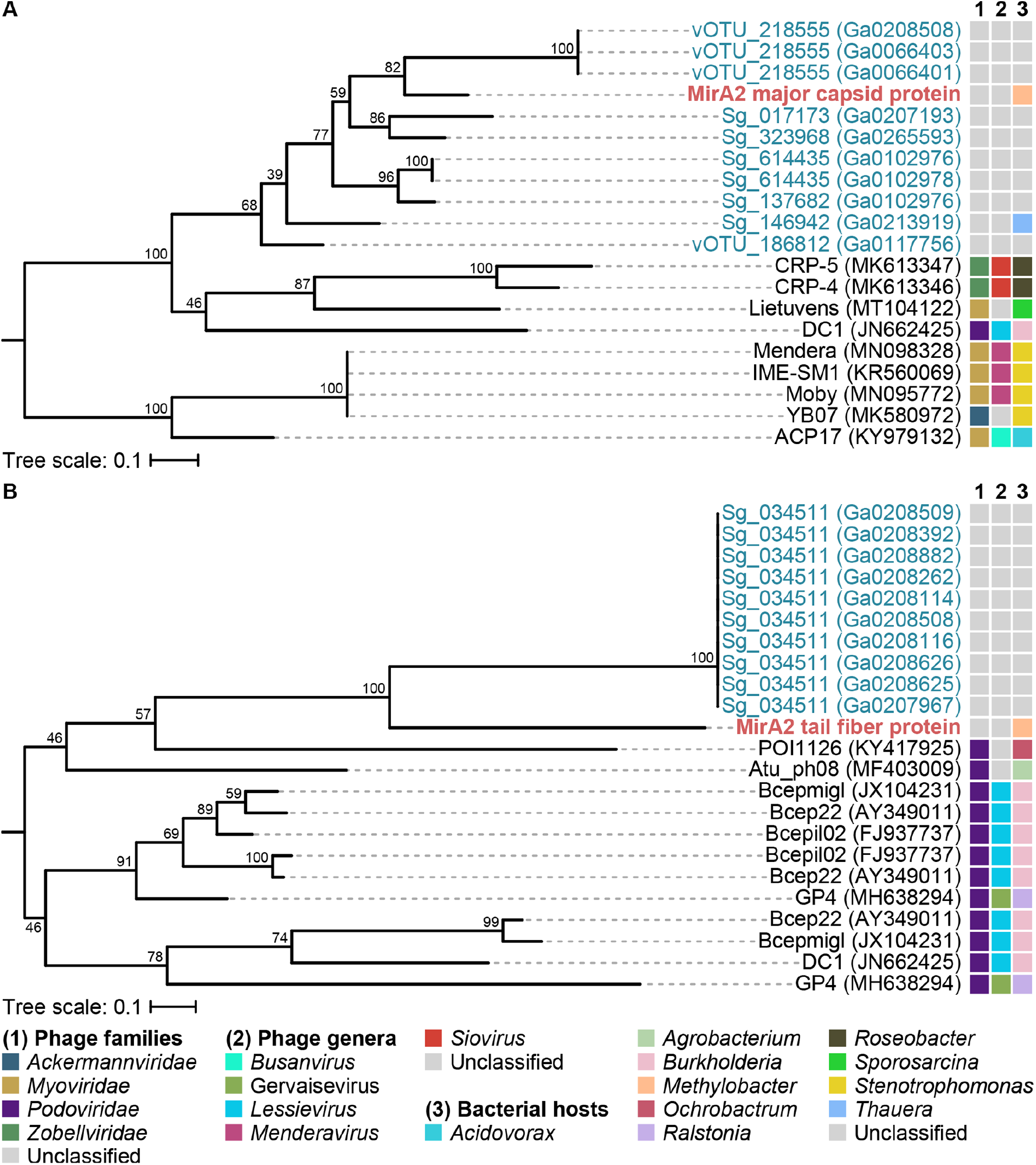
Phylogenetic analysis of induced phage MirA2 major capsid (A) and tail-associated (B) proteins. Midpoint-rooted, maximum-likelihood trees were reconstructed from amino acid sequences (123 and 214 amino acid positions, respectively). Bootstrap support is indicated on the nodes. The scale bar represents amino acid substitutions per site. Tree tip colors indicate genes from the induced phage in this study (red), from NCBI hits (black), or from uncultivated sources (blue). Additional annotations include known (for NCBI hits) or predicted (for IMG/VR hits) 1) phage family designations, 2) phage genus designations, and 3) bacterial hosts.

**Figure 10.**
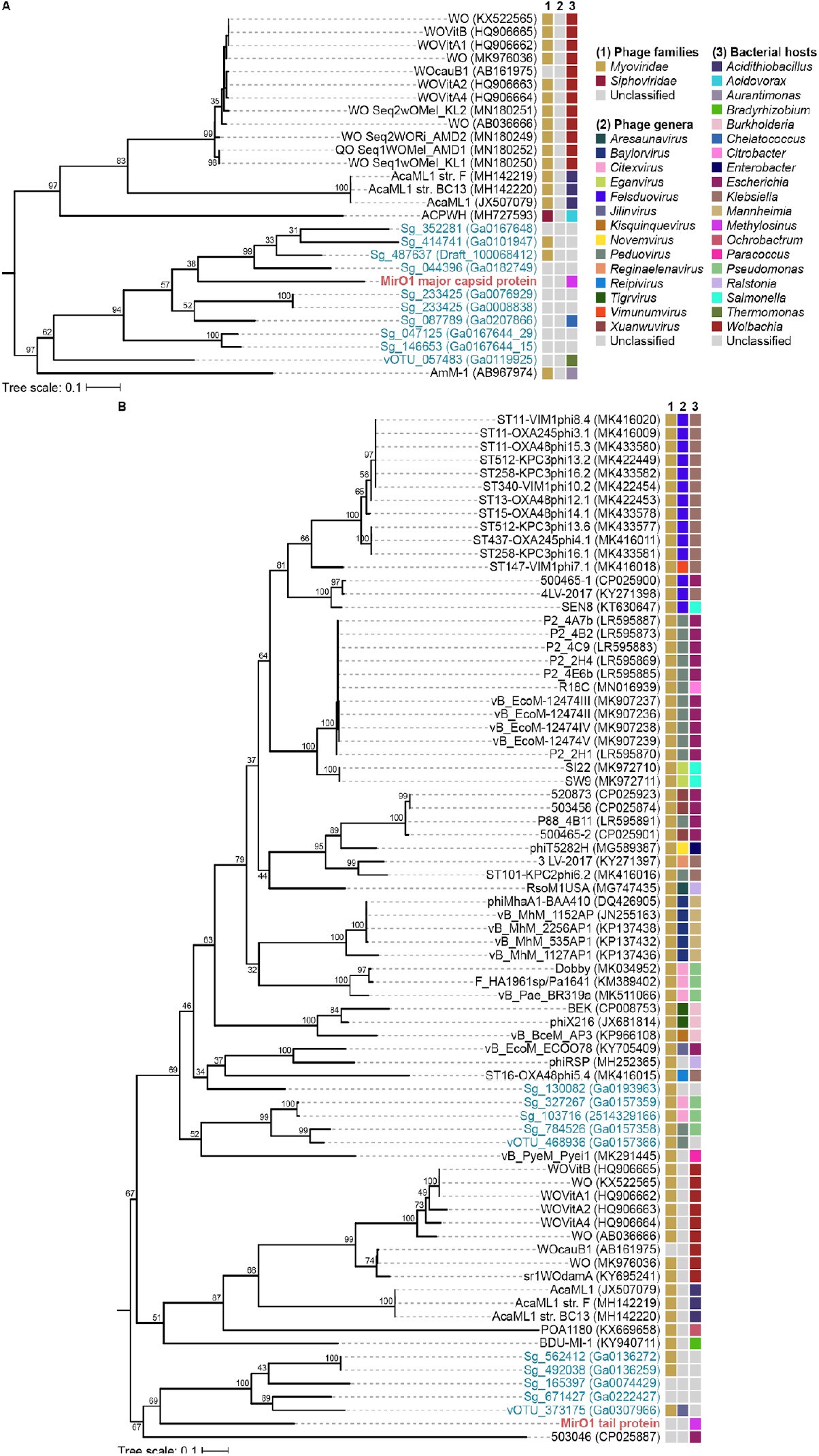
Phylogenetic analysis of induced phage MirO1 major capsid (A) and tail-associated (B) proteins. Midpoint-rooted, maximum-likelihood trees were reconstructed from amino acid sequences (151 and 231 amino acid positions, respectively). Bootstrap support is indicated on the nodes. The scale bar represents amino acid substitutions per site. Tree tip colors indicate genes from the induced phage in this study (red), from NCBI hits (black), or from uncultivated sources (blue). Additional annotations include known (for NCBI hits) or predicted (for IMG/VR hits) 1) phage family designations, 2) phage genus designations, and 3) bacterial hosts.

For all three phages, phylogenetic analysis showed some level of clustering primarily with ‘unclassified’ samples from the environmental (IMG/VR) database, and even less clustering with the top hits from NCBI.

The phylogenetic tree of the major capsid protein of phage MirA1 (Figure 8A) clusters to some degree with unclassified environmental phage samples (Sg_222092, Sg_579391, Sg_491958) that had *Methylosarcina* identified as their putative bacterial host. The lengths of the branches reveal that the capsid protein is even more distantly related to the phage relatives from the NCBI database that belong to the phage family previously designated as *Siphoviridae* with a wide variety of hosts, including *Caulobacter* and *Synechococcus*. The highest patristic distance (PD), calculated from this tree between MirA1 capsid protein and phage CcrColossus (currently designated as a *Siphoviridae*), is 1.871, while the lowest distance of 0.0741 was identified for environmental sample Sg_222092.

The significant distant relatedness was also observed to a greater degree for the major tail protein of MirA1 (Figure 8B) with environmental phage samples (Sg_890835, Sg_817494, Sg_868545), as well as *Stenotrophomonas* targeting phages from the NCBI database. The PD calculated from this tree, between MirA1 tail protein and NCBI and environmental samples, was 1.086 or greater.

Interestingly, the phylogenetic tree analysis of the major capsid protein and major tail protein of MirA1 showed a unique clustering with no overlap between the closest relatives of both trees.

For the other two phages, MirA2 and MirO1, similar trends were seen for the capsid (Figures 9A and 10A) and tail proteins (Figures 9B and 10B). In each case, the PD values ranged from 0.811 to 2.916 for environmental and NCBI samples, respectively.

## Discussion

The general lack of knowledge on methanotroph phage biology is at least partly because of comparatively limited studies outside of the established model organisms or pathogens. To date, few studies characterizing methanotroph phages have been reported. Recent work identified phage sequences associated with methanotrophs from metagenomic and transcriptomic datasets (Gambelli et al., 2016; Chen et al., 2020); and prior to this, two studies from the 1980s presented rudimentary characterization of methanotroph phages isolated from various sources, including cattle rumen and fish gut microbiomes (Tyutikov, Bespalova, Rebentish, Aleksandrushkina, & Krivisky, 1980; Galchenko, & Tikhonenko, 1983). However, the whereabouts and further characterization of these phages remain unclear. Since then, information on phages of methanotrophs has generally been anecdotal. Even less is known about the impacts of phage infections on the population and activity of methanotrophic bacteria. Considering the important role phages play on microbial populations and ecosystem functions, and the significant contributions of methanotrophic bacteria to the global carbon cycle and to advances in biotechnological applications, it is clear that methanotroph phages should not remain in the shadows.

### Prophage induction in methanotrophic bacteria

In the present study we predicted, induced, isolated, and characterized lysogenic phages of methanotrophs. Using MitC, we were able to successfully induce and visualize phages from three out of five strains of methanotrophs tested: *M. marinus* A45, *M. trichosporium* OB3b, and *M. album* BG8, confirming that this method can be used for methanotrophs.

AUC analysis (Figure 1) and statistical tests have shown that MitC inductions were more reliable at a concentration of 2 μg/ml rather than 1 μg/ml. The smaller AUC observed with the higher drug concentration can be correlated to earlier and more effective cell lysis and release of phage virions. This suggests that, at the higher MitC concentration, prophage induction was more rapid and prevalent in the host population, thus accelerating the observed population lysis.

As mentioned above, samples were analyzed under TEM using uranyl acetate staining when ‘symptoms’ of induction were observed during culture growth, including the described reductions in OD and AUC, precipitation and cell clumping. Induced phages visually appeared to be Siphovirus morphotypes all featuring symmetrical polyhedral heads and long tails.

Based on the cell density obtained in the induction experiments, the titer of the resulting lysate was expected to be relatively low. Specific titer numbers could not be confirmed, however, due to the difficulty/impossibility of growing methanotrophs as bacterial lawns on plates and to the presence of the prophages in the bacterial host genomes available conferring a protection against further infection. This highlights a substantial gap in the field of phage research, whereby the majority of protocols are biased towards bacteria which are easily culturable on plates, and to the availability of prophage-less susceptible host species or variants. With 99% of environmental bacteria predicted to be “unculturable” (Sharma, Ranjan, Kapardar, & Grover, 2005) – meaning that current or standard laboratory techniques are not sufficient to enable these microorganisms to grow in lab environments (Stewart, 2012) – new methodologies for the determination of standard phage measurements such as titer and burst size must be developed for methanotrophs.

It should be noted that the addition of MitC to a culture did not always result in the successful identification of an induced phage. For instance, despite PHASTER, Phigaro, PhageBoost and IslandViewer all detecting putative phage regions in both *Methylocystis sp*. Rockwell and *M. denitrificans* FJG1, and both strains showing decreases in OD in response to MitC addition, no phage particles could be identified through TEM analysis. Several factors could be contributing to such results, such as the possibility of MitC killing the bacteria via the SOS response instead of via the induction of a viable phage (Kohanski, Dwyer, Hayete, Lawrence, & Collins, 2007; Miller et al., 2004) and the limit of detection of TEM (which is approximately 10 pfu/ml). Perhaps, for these organisms, different known inducers such as UV radiation or peroxide could be tested. Additionally, the function of many phage-associated genes in the genomes of these bacteria was not fully elucidated (many remain annotated as ‘hypothetical proteins’), hence many regions flagged as potential prophages by different softwares could be unrelated to phages or have degraded over time, losing some of the genes required to re-enter the lytic life cycle (Bobay et al., 2012; Howard-Varona et al., 2017). However, comparative genomics revealed that the uninduced prophages identified displayed high similarities to phages assigned to families previously designated as *Siphoviridae* and *Myoviridae*.

### Prediction of prophages

Since little is known about methanotroph phages they have not been directly used in the development of prophage prediction algorithms. They are thus an interesting system to use for evaluation of these algorithms. Comparative analysis of the predictions from PHASTER, Phigaro, PhageBoost and IslandViewer with the sequences of the induced phages provided valuable information on these useful tools. Firstly, PHASTER and Phigaro rely heavily on well annotated phage genes and other factors such as GC content (Arndt et al., 2016). Considering the current gaps in knowledge in phage genomics – demonstrated by our NCBI BLAST output that was dominated by genes for hypothetical proteins of bacterial hosts (Figures 5-7, Supplementary Figures S3-S5) and the relatively high GC content of the induced phages (Table 1) – this approach may be limited by current information biases. In regions where both PHASTER and Phigaro identified putative prophages (Figure 3, Supplementary Figure S2), the regions themselves usually aligned with moderate to high percent identities. Discrepancies occurred mainly at the ends, highlighting the differences in the algorithms used by the programs to determine the cut offs of putative phage regions. Also, each program did uniquely identify potential prophages that its counterparts did not, accentuating the point that no one program should be used in isolation to analyze prophage regions in a bacterial host strain. PhageBoost applies machine learning to biological features such as amino acid composition, GC content, gene length, and coding sequence direction to predict which genes belong to potential phages versus their host. Given the different nature of the PhageBoost analysis, it is interesting that the identified putative phage regions differed from those identified by PHASTER and Phigaro (Figure 3B and Supplementary Figure S2A). In some instances, the three tools did identify overlapping regions (Figure 3A). Of the three sequenced phages, MirA1 was identified as a potential prophage by both Phigaro and PHASTER (Figure 3A). MirA2 was identified by Phigaro, PhageBoost and IslandViewer (Figure 3A). Interestingly, the sequence of phage MirO1 was not identified by any of the four algorithms. Overall, this analysis emphasizes the gaps in phage biology, including how these entities evolved, how they transfer genes across diverse hosts, and the full extent of their environmental roles.

### Genetic diversity within and between induced prophage genomes

Perhaps the most interesting finding in the present study is the genetic diversity observed between and within the genomes of the induced prophages (Figures 5-10). While these results suggest a prevalence of horizontal gene transfer in methanotroph phage evolution, it is also important to note that since BLAST analysis relies on the pool of genomic information available at any given time, it is also reliant on the relatively small amount of genomic information related to methanotrophic bacteria and phages currently available in databases. For example, while there are over 23,000 *Escherichia coli* genomes in the NCBI database and 323 coliphage genomes in the *Caudovirales* database, only 651 gammaproteobacterial methanotroph genomes are publicly accessible at this time.

Phylogenetic analysis based on whole phage genomes (Supplementary Figures 6-8), major capsid proteins, and structural tail-associated proteins (major tail proteins and tail fiber proteins) (Figures 8-10) was used to place each of the induced prophages within the *Caudovirales* order and assess their genetic diversity. Analysis against the complete NCBI database, in addition to the *Caudovirales* database, was included to address the possibility that the induced methanotroph phages would be most closely related to not-yet-identified prophages. Despite this wide-ranging net, BLAST analysis revealed low percentage identity values across both database screens for all induced methanotroph phages (Figures 5-7), suggesting sequence uniqueness.

Focusing on the individual methanotroph phages, nBLAST and pBLAST of phage MirA1 against the *Caudovirales* database suggested its closest relatives were classified as part of the phage family previously designated as *Siphoviridae* (Figure 5D, 5F and Figure 8); also confirmed by the morphology of the phage (Figure 2A, 2B). Phage MirA2 revealed a phylogeny with closest relatives classified as part of the previously defined *Podoviridae* and *Myoviridae* families (Figure 9). For phage MirO1, phylogenetic analyses placed its major capsid protein gene and tail fiber gene found closest relatives previously classified within the *Myoviridae* family (Figure 10), whereas nBLAST analysis of its full genome identified the closest relatives previously classified as part of the *Siphoviridae* and *Myoviridae* families (Figure 7D, 7F). TEM analysis of this phage showed a Siphovirus morphology (Figure 2E, 2F). It is important to note that the *Siphoviridae, Myoviridae*, and *Podoviridae* family designations, which were historically based on morphology, are currently being revisited based on genomic diversity, and are now considered to each comprise numerous phage families (Turner et al. 2021). In fact, there is a recognized disconnect between phage morphology and genome sequence analysis – a concept that complicates phage classification using the classical approach based solely on morphology (Adriaenssens et al., 2015; Lavigne et al., 2009; Lavigne et al., 2008). In fact, overall, BLAST analysis against the *Caudovirales* database and phylogenetic analysis demonstrated that phage family identification was not tied to high percentage identities, with an average percentage identity of 32.3% (Figures 5-7). Another example includes the analysis of the major capsid proteins of phages MirA2 and MirO1. In both instances, the major capsid proteins clustered within more than one previously designated phage family. This pervasive genetic mosaicism was most likely caused by horizontal gene transfer (Botstein, 1980; Brüssow & Desiere, 2001; Juhala et al., 2000; G. H. Wang et al., 2016) along with limited genomic information on sequences of methanotroph phages. A signature of potential gene transfer within phage families was seen in both MirA2 and MirO1 phages.

These findings highlight the fact that phages MirA1, MirA2, and MirO1 are each representative of new phage species, genera and families. As such, we propose MirA1 as a representative of the new phage family, genus and species “*Britesideviridae inducovirus miraone*”, MirA2 as a representative of “*Patronusviridae enigmavirus miratwo*”, and MirO1 as a representative of “*Kainiviridae tripudiumvirus miroone*”.

In performing nBLAST searches of the complete phage genomes against the whole NCBI database, it was thought that the phage genomes or gene sequences would be similar to those found in other methanotrophic bacteria (Figures 5-7). For instance, pBLAST analysis of genes from phage MirO1 led to a few top hits associated with genomes of methanotrophs (Suppl. Fig. S5). Its DNA primase showed the highest identity with a gene product from the genome of *Methylosinus* sp. 3S-1, while its portal protein and baseplate assembly protein had the highest identity with sequences from the genome of *Methylosinus* sp. PW1. The top hits for the portal protein gene also included a sequence from *Methylocella tundrae*. But surprisingly, except in a few cases, the top five hits with highest identity for all three phages were unrelated to methanotrophs. No methanotroph-related sequence ranked in the top five hits against the genome or individual genes of phage MirA1 (Figure 5 and Suppl. Fig. S3). Similar analysis for phage MirA2 led to one identification of a methanotroph phage capsid protein sequence from *Methylophilales* (Figure 6D). It should be noted that this *Methylophilales* phage sequence had been inferred from metagenomic analysis and that the potential phage was not cultured.

In addition to providing valuable information on the origin and potential evolutionary pathway of the given phage genomes and proteins investigated, the genetic analyses performed in this study established a framework for the breadth of diversity of methanotroph phages. Again, the fact that the most closely identified relatives are not phages of methanotrophic bacteria, or even related strains, suggests that the databases used (both microbial and phage-based) contain limited information on these entities and have a bias towards the much more heavily studied health-associated bacterial strains.

## Conclusion

Although little is known about phages of methanotrophs, early evidence suggests they are diverse and abundant. In this study, four software programs, PHASTER, Phigaro, PhageBoost and IslandViewer, were used to identify putative prophage regions in five methanotrophic bacteria. The analyses showed that the four algorithms are not always in agreement with regards to what constitutes a putative prophage region, and that the ends of the phage regions differ when putative prophage regions overlap. MitC was used to attempt to induce lysogenized phages from the five methanotrophs. In total four prophages were induced (two from *M. marinus* A45, one from *M. album* BG8, and one from *M. trichosporium* OB3b), three of which were imaged by TEM and subsequently sequenced (MirA1, MirA2, MirO1).

Phylogenetic and BLAST analyses were used to establish the genetic diversity and potential evolutionary history of the induced methanotroph phages. These analyses highlighted the broad diversity and gaps in genetic information of methanotroph phages. This report lays the groundwork for a better understanding of the ecology and development of technologies based on methanotrophic bacteria and their associated phages.

## Acknowledgements

Special thanks to Arlene Oatway for her technical support and guidance in visualizing induced phage under TEM microscopy and Kevin Jones for bioinformatic support with Phigaro analysis.

Funding was provided by the Canada First Research Excellence Fund – Future Energy Systems, the Alberta Innovates BioSolutions Biofuture program, the Natural Sciences and Engineering Research Council of Canada Discovery grant program, the Canadian Foundation for Innovation – John R. Evans Leaders Fund, the Government of Alberta – Ministry of Economic Development and Trade Small Equipment Grant, and the Alberta Innovates Accelerating Innovations into CarE program.

## Notes

### Competing Interest Statement

The authors have declared no competing interest.

## References

Angly, F. E., Felts, B., Breitbart, M., Salamon, P., Edwards, R. A., Carlson, C., … Rohwer, F. (2006). The marine viromes of four oceanic regions. PLoS Biology, 4(11), e368. https://doi.org/10.1371/journal.pbio.0040368

Arndt, D., Grant, J. R., Marcu, A., Sajed, T., Pon, A., Liang, Y., & Wishart, D. S. (2016). PHASTER: a better, faster version of the PHAST phage search tool. Nucleic Acids Research, 44(W1), W16–W21. https://doi.org/10.1093/nar/gkw387

Bertelli, C., Laird, M. R., Williams, K. P., Lau, B. Y., Hoad, G., Winsor, G. L., & Brinkman, F. S. (2017). IslandViewer 4: expanded prediction of genomic islands for larger-scale datasets. Nucleic Acids Research, 45(W1), W30–W35. https://doi.org/10.1093/nar/gkx343

Bobay, L.-M., Rocha, E. P. C., & Touchon, M. (2012). The adaptation of temperate bacteriophages to their host genomes. Molecular Biology and Evolution, 30(4), 737–751. https://doi.org/10.1093/molbev/mss279

Bortolaia, V., Kaas, R. S., Ruppe, E., Roberts, M. C., Schwarz, S., Cattoir, V., … Aarestrup, F. M. (2020). ResFinder 4.0 for predictions of phenotypes from genotypes. Journal of Antimicrobial Chemotherapy. https://doi.org/10.1093/jac/dkaa345

Botstein, D. (1980). A theory of modular evolution for bacteriophages. Annals of the New York Academy of Sciences, 354(1), 484–491. https://doi.org/10.1111/j.1749-6632.1980.tb27987.x

Brüssow, H., & Desiere, F. (2001). Comparative phage genomics and the evolution of Siphoviridae: Insights from dairy phages. Molecular Microbiology, 39(2), 213–223. https://doi.org/10.1046/j.1365-2958.2001.02228.x

Cantera, S., Muñoz, R., Lebrero, R., López, J. C., Rodríguez, Y., & García-Encina, P. A. (2018). Technologies for the bioconversion of methane into more valuable products. Current Opinion in Biotechnology, 50, 128–135. https://doi.org/10.1016/j.copbio.2017.12.021

Casjens, S. R., & Hendrix, R. W. (2015). Bacteriophage lambda: Early pioneer and still relevant. Virology, 479–480, 310–330. https://doi.org/10.1016/j.virol.2015.02.010

Chen, L. X., Méheust, R., Crits-Christoph, A., McMahon, K. D., Nelson, T. C., Slater, G. F., … Banfield, J. F. (2020). Large freshwater phages with the potential to augment aerobic methane oxidation. Nature Microbiology, 5, 1504–1515. https://doi.org/10.1038/s41564-020-0779-9

Cook, R., Brown, N., Redgwell, T., Rhitman, B., Barnes, M., Clokie, M., Stekel, D.J., Hobman, J.L., Jones, M.A., & Millard, A. (2021) INfrastructure for a PHAge REference Database: Identification of large-scale biases in the current collection of phage genomes. PHAGE: Therapy, Applications and Research, 2(4), 214–222. https://doi.org/10.1101/2021.05.01.442102.

De Paepe, M., Tournier, L., Moncaut, E., Son, O., Langella, P., & Petit, M.-A. (2016). Carriage of lambda latent virus is costly for its bacterial host due to frequent reactivation in monoxenic mouse intestine. PLoS Genetics, 12(2), e1005861. https://doi.org/10.1371/journal.pgen.1005861

Eddy, S. R. (2011). Accelerated profile HMM searches. PLoS Computational Biology, 7(10), e1002195. https://doi.org/10.1371/journal.pcbi.1002195

Edgar, R. C. (2004). MUSCLE: a multiple sequence alignment method with reduced time and space complexity. BMC Bioinformatics, 5, 113. https://doi.org/10.1186/1471-2105-5-113

Emerson, J. B., Roux, S., Brum, J. R., Bolduc, B., Woodcroft, B. J., Jang, H. Bin, … Sullivan, M. B. (2018). Host-linked soil viral ecology along a permafrost thaw gradient. Nature Microbiology, 3, 870–880. https://doi.org/10.1038/s41564-018-0190-y

Gambelli, L., Cremers, G., Mesman, R., Guerrero, S., Dutilh, B. E., Jetten, M. S. M., … van Niftrik, L. (2016). Ultrastructure and viral metagenome of bacteriophages from an anaerobic methane oxidizing Methylomirabilis bioreactor enrichment culture. Frontiers in Microbiology, 7, 1740. https://doi.org/10.3389/fmicb.2016.01740

Grazziotin, A. L., Koonin, E. V., & Kristensen, D. M. (2017). Prokaryotic Virus Orthologous Groups (pVOGs): A resource for comparative genomics and protein family annotation. Nucleic Acids Research, 45(D1), D491–D498. https://doi.org/10.1093/nar/gkw975

Hare, J. M., Ferrell, J. C., Witkowski, T. A., & Grice, A. N. (2014). Prophage induction and differential RecA and UmuDAb transcriptome regulation in the DNA damage responses of Acinetobacter baumannii and Acinetobacter baylyi. PloS One, 9(4), e93861–e93861. https://doi.org/10.1371/journal.pone.0093861

Henard, C. A., Smith, H. K., & Guarnieri, M. T. (2017). Phosphoketolase overexpression increases biomass and lipid yield from methane in an obligate methanotrophic biocatalyst. Metabolic Engineering, 41, 152–158. https://doi.org/10.1016/j.ymben.2017.03.007

Howard-Varona, C., Hargreaves, K. R., Abedon, S. T., & Sullivan, M. B. (2017). Lysogeny in nature: mechanisms, impact and ecology of temperate phages. The ISME Journal, 11(7), 1511–1520. https://doi.org/10.1038/ismej.2017.16

Jiang, H., Chen, Y., Jiang, P., Zhang, C., Smith, T. J., Murrell, J. C., & Xing, X. H. (2010). Methanotrophs: Multifunctional bacteria with promising applications in environmental bioengineering. Biochemical Engineering Journal, 49(3), 277–288. https://doi.org/10.1016/j.bej.2010.01.003

Juhala, R. J., Ford, M. E., Duda, R. L., Youlton, A., Hatfull, G. F., & Hendrix, R. W. (2000). Genomic sequences of bacteriophages HK97 and HK022: Pervasive genetic mosaicism in the lambdoid bacteriophages. Journal of Molecular Biology, 299(1), 27–51. https://doi.org/10.1006/jmbi.2000.3729

Kaserer, H. (1905). Uber die oxidation des wasserstoffes und des methan durch mikroorganismen. (The oxidation of hydrogen and methane by microorganisms.) Z. Landw. Versuchsw. in Osterreich, 8, 789–792.

Katoh, K., & Toh, H. (2008). Recent developments in the MAFFT multiple sequence alignment program. Briefings in Bioinformatics, 9(4), 286–298. https://doi.org/10.1093/bib/bbn013

Kearse, M., Moir, R., Wilson, A., Stones-Havas, S., Cheung, M., Sturrock, S., … Drummond, A. (2012). Geneious Basic: An integrated and extendable desktop software platform for the organization and analysis of sequence data. Bioinformatics. https://doi.org/10.1093/bioinformatics/bts199

Khmelenina, V. N., Kalyuzhnaya, M. G., Sakharovsky, V. G., Snzina, N. E., Trotsenko, Y. A., & Gottschalk, G. (1999). Osmoadaptation in halophilic and alkaliphilic methanotrophs. Archives of Microbiology, 172, 321–329. https://doi.org/10.1007/s002030050786

Kilic, A. O., Pavlova, S. I., Ma, W. G., & Tao, L. (1996). Analysis of Lactobacillus phages and bacteriocins in American dairy products and characterization of a phage isolated from yogurt. Applied and Environmental Microbiology, 62(6), 2111–2116.

Kleinheinz, K. A., Joensen, K. G., & Larsen, M. V. (2014). Applying the ResFinder and VirulenceFinder web-services for easy identification of acquired antibiotic resistance and E. coli virulence genes in bacteriophage and prophage nucleotide sequences. Bacteriophage, 4, e27943. https://doi.org/10.4161/bact.27943

Lee, O. K., Hur, D. H., Nguyen, D. T. N., & Lee, E. Y. (2016). Metabolic engineering of methanotrophs and its application to production of chemicals and biofuels from methane. Biofuels, Bioproducts and Biorefining, 10(6), 848–863. https://doi.org/10.1002/bbb.1678

Lee, S., Sieradzki, E. T., Nicolas, A. M., Walker, R. L., Firestone, M. K., Hazard, C., Nicol, G. W. (2021) Methane-derived carbon flows into host-virus networks at different trophic levels in soil. Proceedings of the National Academy of Sciences of the United States of America. 118(32), e2105124118. https://doi.org/10.1073/pnas.2105124118.

Letunic, I., & Bork, P. (2021). Interactive Tree Of Life (iTOL) v5: an online tool for phylogenetic tree display and annotation. Nucleic Acids Research, 49(W1), W293–296. https://doi.org/10.1093/nar/gkab301

Maslowska, K. H., Makiela-Dzbenska, K., & Fijalkowska, I. J. (2019). The SOS system: A complex and tightly regulated response to DNA damage. Environmental and Molecular Mutagenesis, 60(4), 368–384. https://doi.org/10.1002/em.22267

McDonald, J. E., Smith, D. L., Fogg, P. C. M., McCarthy, A. J., & Allison, H. E. (2010). High-throughput method for rapid induction of prophages from lysogens and its application in the study of Shiga toxin-encoding Escherichia coli strains. Applied and Environmental Microbiology, 76(7), 2360–2365. https://doi.org/10.1128/AEM.02923-09

Nishimura, Y., Yoshida, T., Kuronishi, M., Uehara, H., Ogata, H., & Goto, S. (2017) ViPTree: the viral proteomic tree server. Bioinformatics, 33(15), 2379–2380. https://doi.org/10.1093/bioinformatics/btx157

Nguyen, L. T., Schmidt, H. A., Von Haeseler, A., & Minh, B. Q. (2015). IQ-TREE: A fast and effective stochastic algorithm for estimating maximum-likelihood phylogenies. Molecular Biology and Evolution, 32(1), 268–274. https://doi.org/10.1093/molbev/msu300

Otsuji, N., Sekiguchi, M., Iijima, T., & Takagi, Y. (1959). Induction of phage formation in the lysogenic Escherichia coli K-12 by Mitomycin C. Nature, 184(4692), 1079–1080. https://doi.org/10.1038/1841079b0

Posada, D., & Crandall, K. A. (1998). MODELTEST: Testing the model of DNA substitution. Bioinformatics, 14(9), 817–818. https://doi.org/10.1093/bioinformatics/14.9.817

Puapermpoonsiri, U., Ford, S. J., & van der Walle, C. F. (2010). Stabilization of bacteriophage during freeze drying. International Journal of Pharmaceutics, 389(1-2), 168–175. https://doi.org/10.1016/j.ijpharm.2010.01.034

Ramisetty, B. C. M., & Sudhakari, P. A. (2019). Bacterial ‘grounded’ prophages: Hotspots for genetic renovation and innovation. Frontiers in Genetics, 10, 65. https://www.frontiersin.org/article/10.3389/fgene.2019.00065

Roskams, J., & Rodgers, L. (eds.) (2002) Lab Ref, Volume 1: A handbook of recipes reagents, and other reference tools for use at the bench. Cold Spring Harbour, CSHL Press.

Roux, S., Páez-Espino, D., Chen, I.-M. A., Palaniappan, K., Ratner, A., Chu, K., Reddy, T. B. K., Nayfach, S., Schulz, F., Call, L., Neches, R. Y., Woyke, T., Ivanova, N. N., Eloe-Fadrosh, E. A., & Kyrpides, N. C. (2021). IMG/VR v3: an integrated ecological and evolutionary framework for interrogating genomes of uncultivated viruses. Nucleic Acid Research, 49(D1), D764-D775. https://doi.org/10.1093/nar/gkaa946

Seemann, T. (2014). Prokka: Rapid prokaryotic genome annotation. Bioinformatics, 30(14), 2068–2069. https://doi.org/10.1093/bioinformatics/btu153

Sharma, R., Ranjan, R., Kapardar, R. K., & Grover, A. (2005). “Unculturable” bacterial diversity: An untapped resource. Current Science, 89(1), 72–77. http://www.jstor.org/stable/24110433

Shiba, S., Terawaki, A., Taguchi, T., & Kawamata, J. (1959). Selective inhibition of formation of deoxyribonucleic acid in Escherichia coli by Mitomycin C. Nature, 183(4667), 1056–1057. https://doi.org/10.1038/1831056a0

Söhngen, N. (1906). Uber bakterien welche methan ab kohlenstoffnahrung und energiequelle gerbrauchen (On bacteria which use methane as a carbon and energy source). Z. Bakteriol. Parazitenk, 15, 513–517.

Stamatakis, A. (2014). RAxML version 8: A tool for phylogenetic analysis and post-analysis of large phylogenies. Bioinformatics, 30(9), 1312–1313. https://doi.org/10.1093/bioinformatics/btu033

Starikova, E. V, Tikhonova, P. O., Prianichnikov, N. A., Rands, C. M., Zdobnov, E. M., & Govorun, V. M. (2019). Phigaro: high throughput prophage sequence annotation. Bioinformatics, 36(12), 3882–3884. https://doi.org/10.1101/598243

Stewart, E. J. (2012). Growing unculturable bacteria. Journal of Bacteriology, 194(16), 4151 LP – 4160. https://doi.org/10.1128/JB.00345-12

Strong, P. J., Xie, S., & Clarke, W. P. (2015). Methane as a resource: Can the methanotrophs add value? Environmental Science & Technology, 49(7), 4001–4018. https://doi.org/10.1021/es504242n

Susskind, M. M., Botstein, D., & Wright, A. (1974). Superinfection exclusion by P22 prophage in lysogens of Salmonella typhimurium: III. Failure of superinfecting phage DNA to enter sieA+ lysogens. Virology, 62(2), 350–366. https://doi.org/https://doi.org/10.1016/0042-s6822(74)90398-5

Tetzschner, A. M. M., Johnson, J. R., Johnston, B. D., Lund, O., & Scheutz, F. (2020). In Silico genotyping of Escherichia coli isolates for extraintestinal virulence genes by use of whole-genome sequencing data. Journal of Clinical Microbiology, 58(10), e01269–20. https://doi.org/10.1128/JCM.01269-20

Turner, D., Kropinski, A.M., & Adriaenssens, E.M. (2021) A Roadmap for Genome-Based Phage Taxonomy. Viruses, 13(3), 506. https://doi.org/10.3390/v13030506

Tyutikov, F. M., Bespalova, I. A., Rebentish, B. A., Aleksandrushkina, N. N., & Krivisky, A. S. (1980). Bacteriophages of methanotrophic bacteria. Journal of Bacteriology, 144(1), 375– 381. Retrieved from https://www.ncbi.nlm.nih.gov/pubmed/6774962

Tyutikov, F. M., Yesipova, V. V, Rebentish, B. A., Bespalova, I. A., Alexandrushkina, N. I., Galchenko, V. V, & Tikhonenko, A. S. (1983). Bacteriophages of methanotrophs isolated from fish. Applied and Environmental Microbiology, 46(4), 917–924.

Wang, G. H., Sun, B. F., Xiong, T. L., Wang, Y. K., Murfin, K. E., Xiao, J. H., & Huang, D. W. (2016). Bacteriophage WO can mediate horizontal gene transfer in endosymbiotic Wolbachia genomes. Frontiers in Microbiology, 7, 1867. https://doi.org/10.3389/fmicb.2016.01867

Wang, X., & Wood, T. K. (2016). Cryptic prophages as targets for drug development. Drug Resistance Updates, 27, 30–38. https://doi.org/10.1016/J.DRUP.2016.06.001

Whittenbury, R., Phillips, K. C., & Wilkinson, J. F. (1970). Enrichment, isolation and some properties of methane-utilizing bacteria. Journal of General Microbiology, 61(2), 205–218. https://doi.org/10.1099/00221287-61-2-205

Zhou, Y., Liang, Y., Lynch, K. H., Dennis, J. J., & Wishart, D. S. (2011). PHAST: a fast phage search tool. Nucleic Acids Research, 39, W347–52. https://doi.org/10.1093/nar/gkr485

