## Supplemental figures for "Lysogenized phages of methanotrophic bacteria show a broad and untapped genetic diversity"

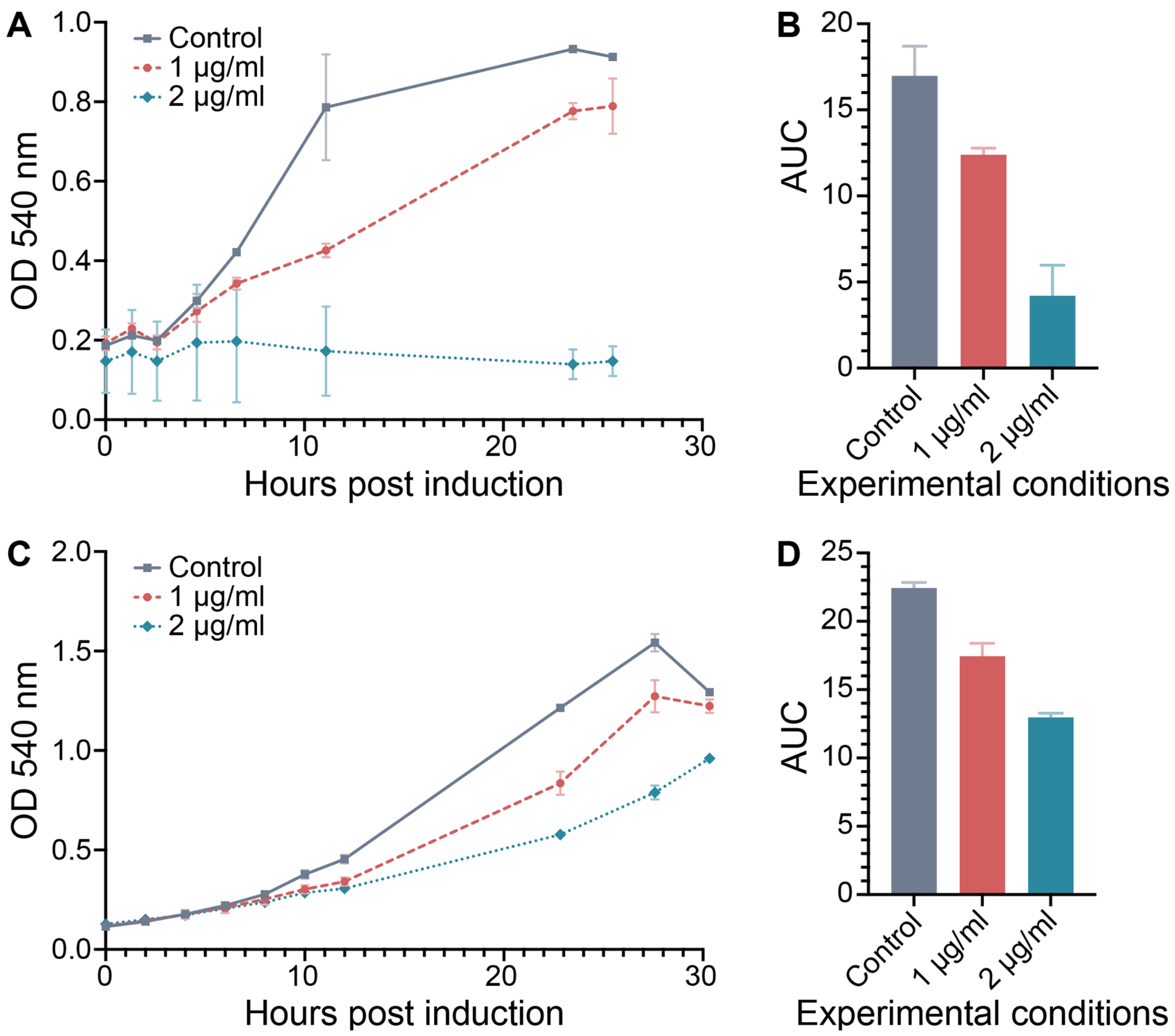


**Supplementary Figure 1.** Induction of potential prophages by addition of mitomycin C to cultures of different methanotrophs. Optical density ($OD_{540}$) and area under the curve of $OD_{540}$ (AUC) are shown for *M. denitrificans* FJG1 (A, B) and *Methylocystis* sp. Rockwell (C, D) after the addition of mitomycin C to concentrations of 1 µg/ml and 2 µg/ml and in control experiments. Error bars denote standard deviation based on n = 3.

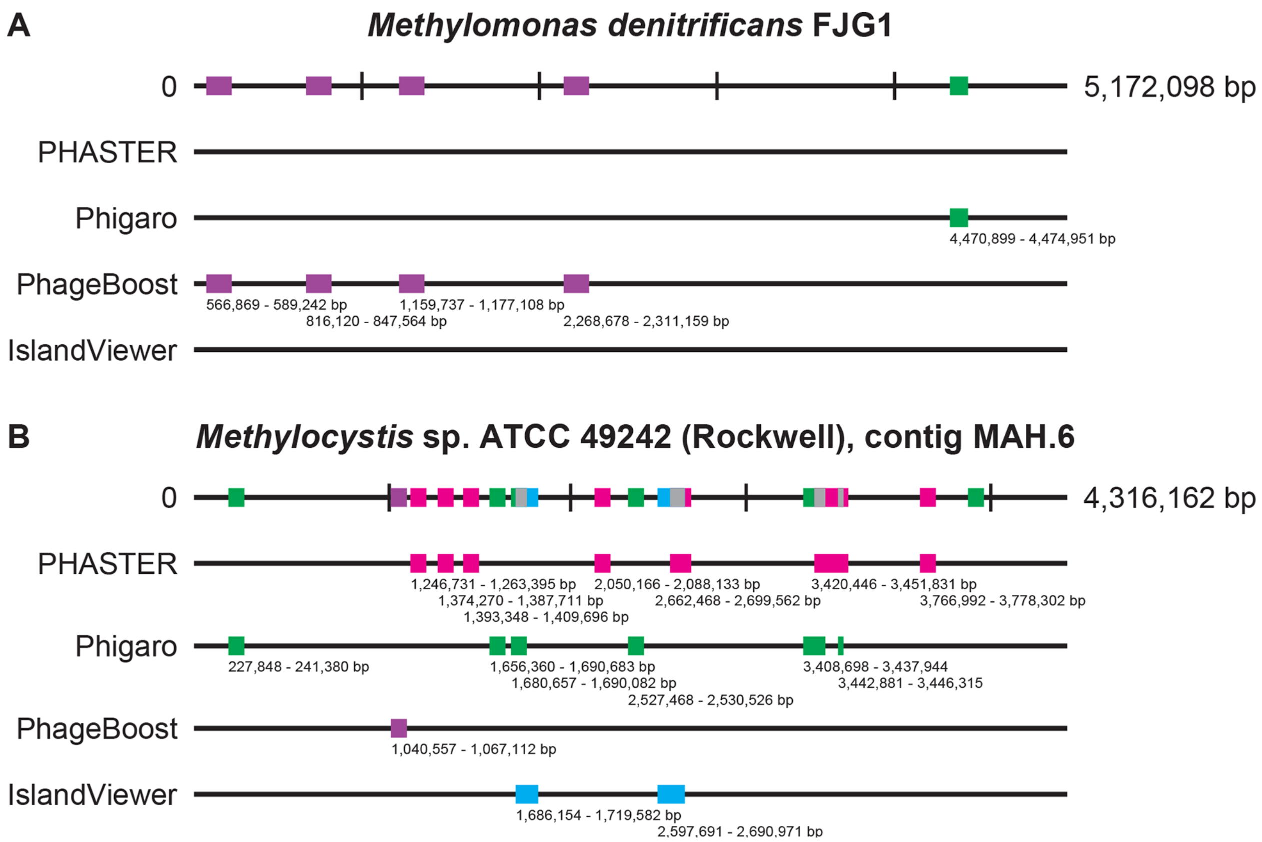


**Supplementary Figure 2.** Computational predictions of prophage regions using PHASTER, Phigaro, PhageBoost, and IslandViewer in the alphaproteobacterial methanotrophs *M. denitrificans* FJG1 (A) and *Methylocystis* sp. Rockwell (B). Reference genome sequences are reported at the top of the figures, followed by visualizations of putative prophage regions identified by PHASTER (magenta), Phigaro (green), PhageBoost (violet), and IslandViewer (blue). Overlapping putative regions are illustrated in grey in the host genome when two predicted regions overlap. The location of each region of interest is indicated by the base pair numbers corresponding to their position in the host genome.

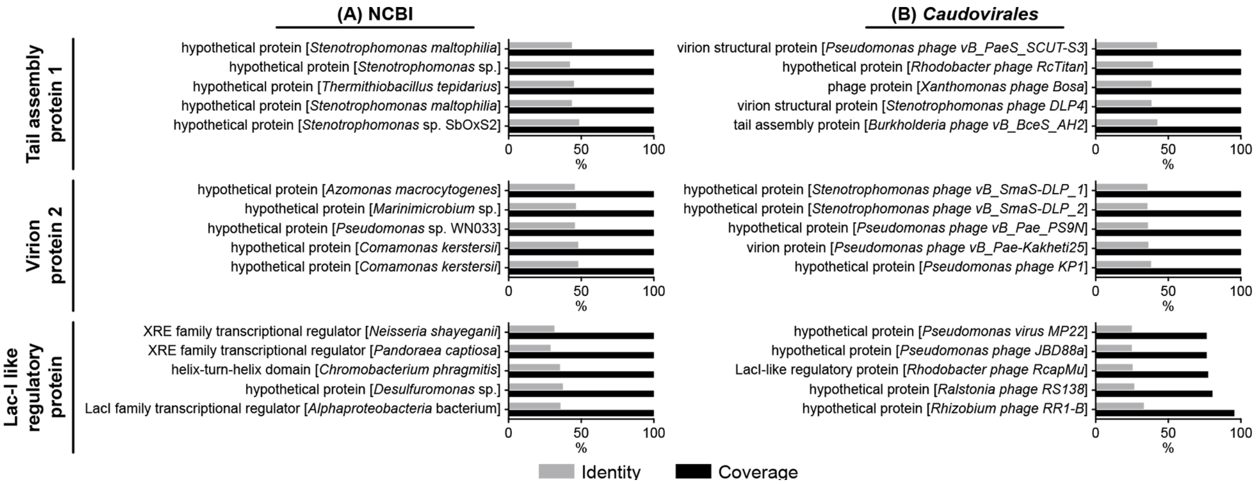


**Supplementary Figure S3.** BLAST analysis of tail assembly protein 1, virion protein 2, and Lac I-like regulatory protein of phage MirA1 against the entire NCBI (A) and *Caudovirales*-only (B) databases. Percent identity (gray) and percent coverage (black) are reported. The top five hits based on percent identity were ordered based on percent coverage.

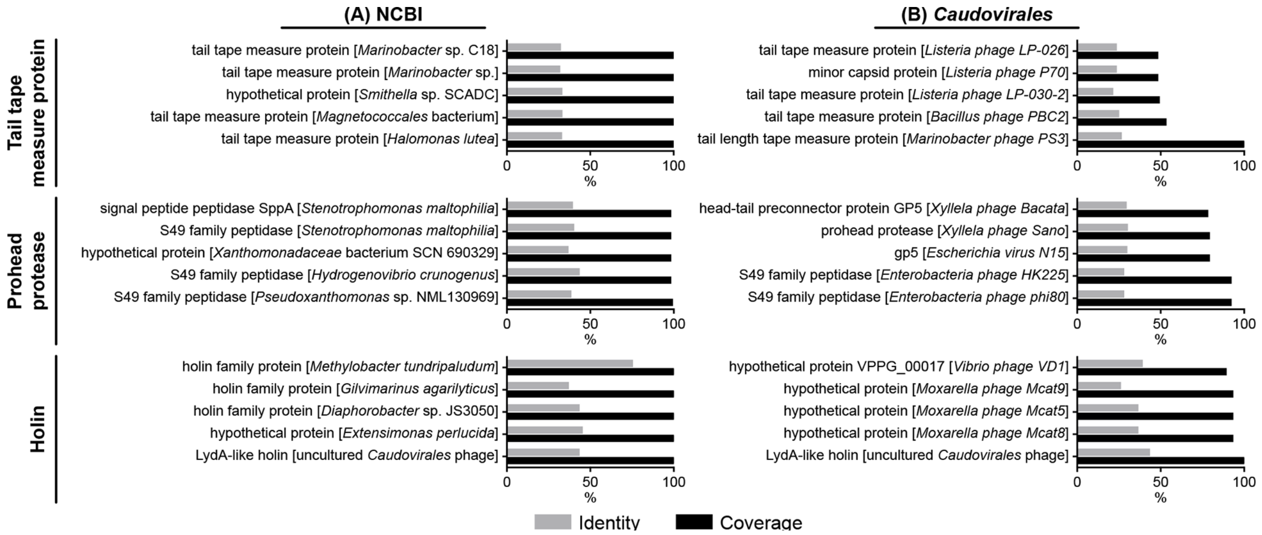


**Supplementary Figure S4.** BLAST analysis of phage tail tape measure protein, prohead protease, and holin of phage MirA2 against the entire NCBI (A) and *Caudovirales*-only (B) databases. Percent identity (gray) and percent coverage (black) are reported. The top five hits based on percent identity were ordered based on percent coverage.

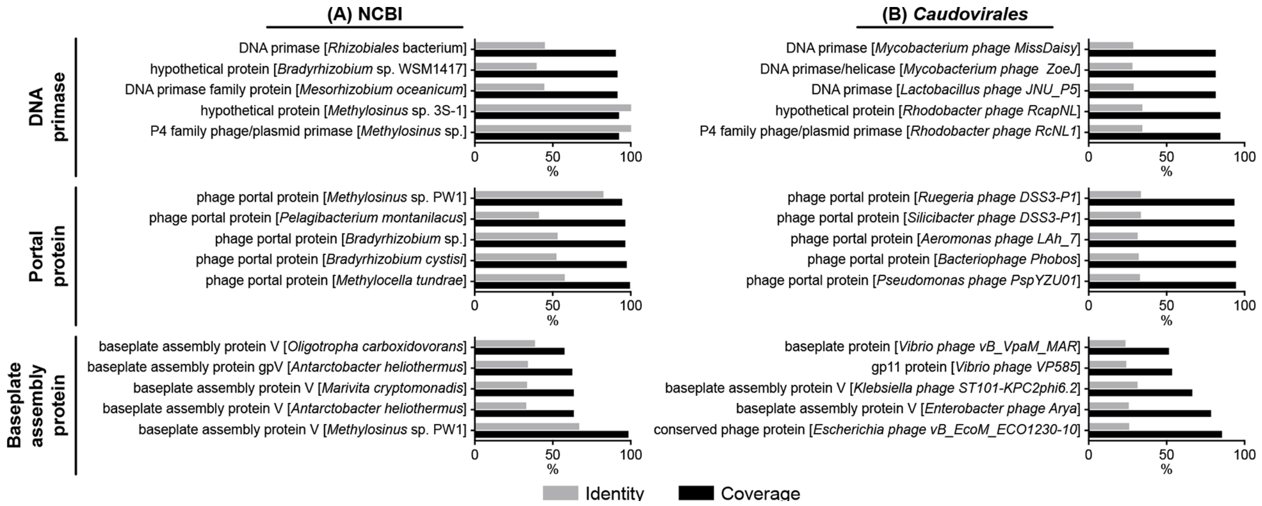


**Supplementary Figure S5.** BLAST analysis of DNA primase, portal protein, and baseplate assembly protein of phage MirO1 against the entire NCBI (A) and *Caudovirales*-only (B) databases. Percent identity (gray) and percent coverage (black) are reported. The top five hits based on percent identity were ordered based on percent coverage.

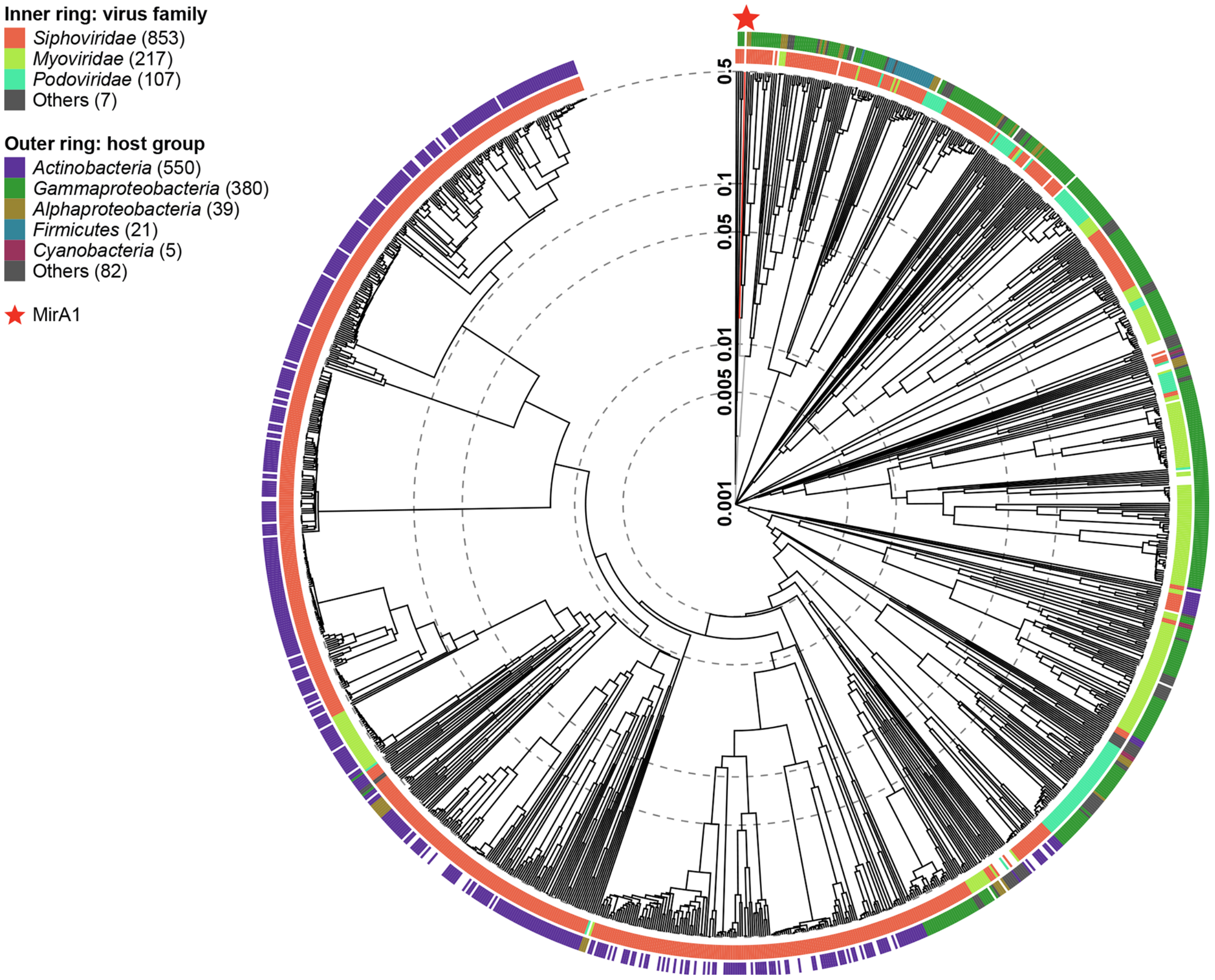


**Supplementary Figure S6.**

Phylogenetic analysis of the entire induced phage MirA1 genome.

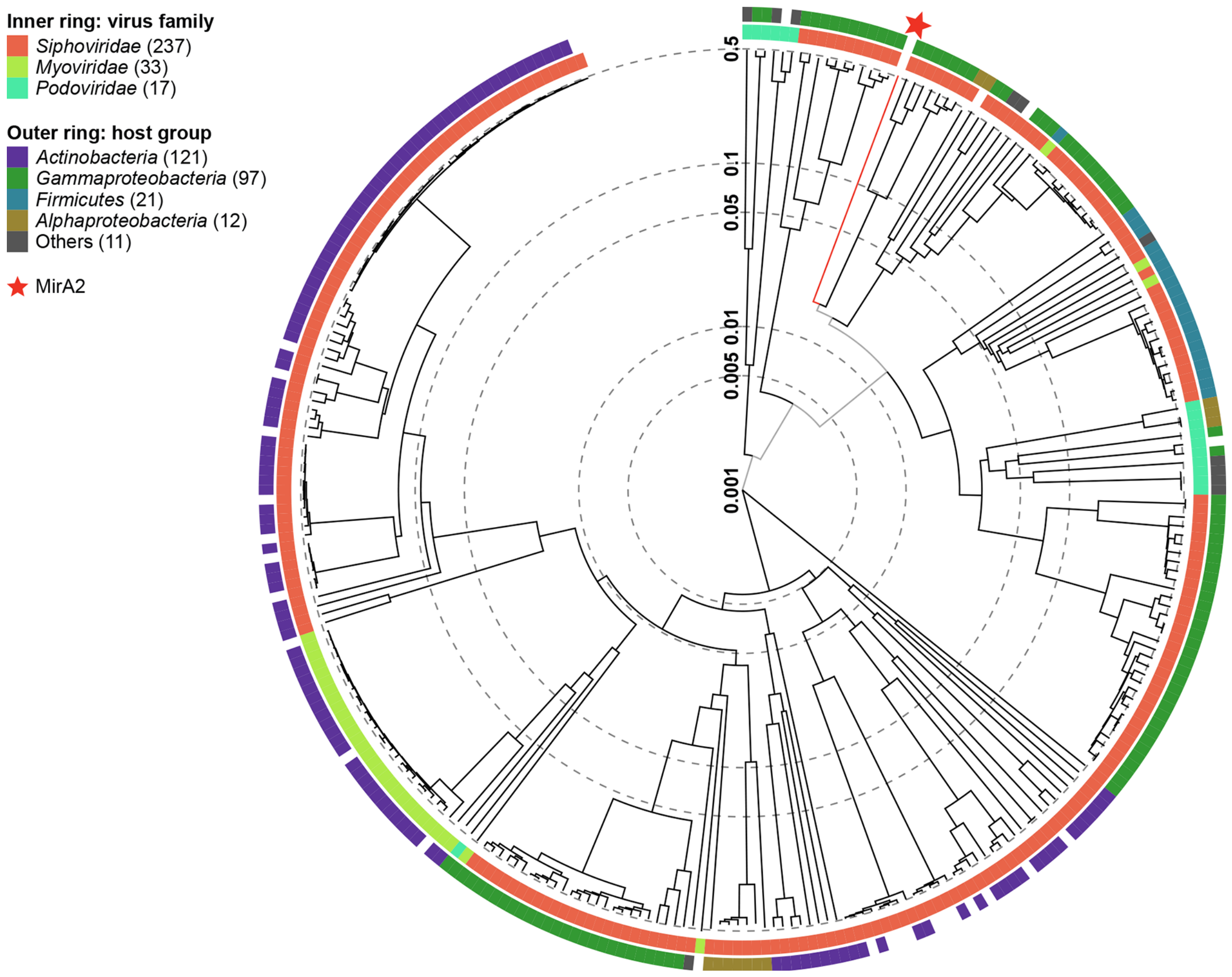


**Supplementary Figure S7.**
Phylogenetic analysis of the entire induced phage MirA2 genome.

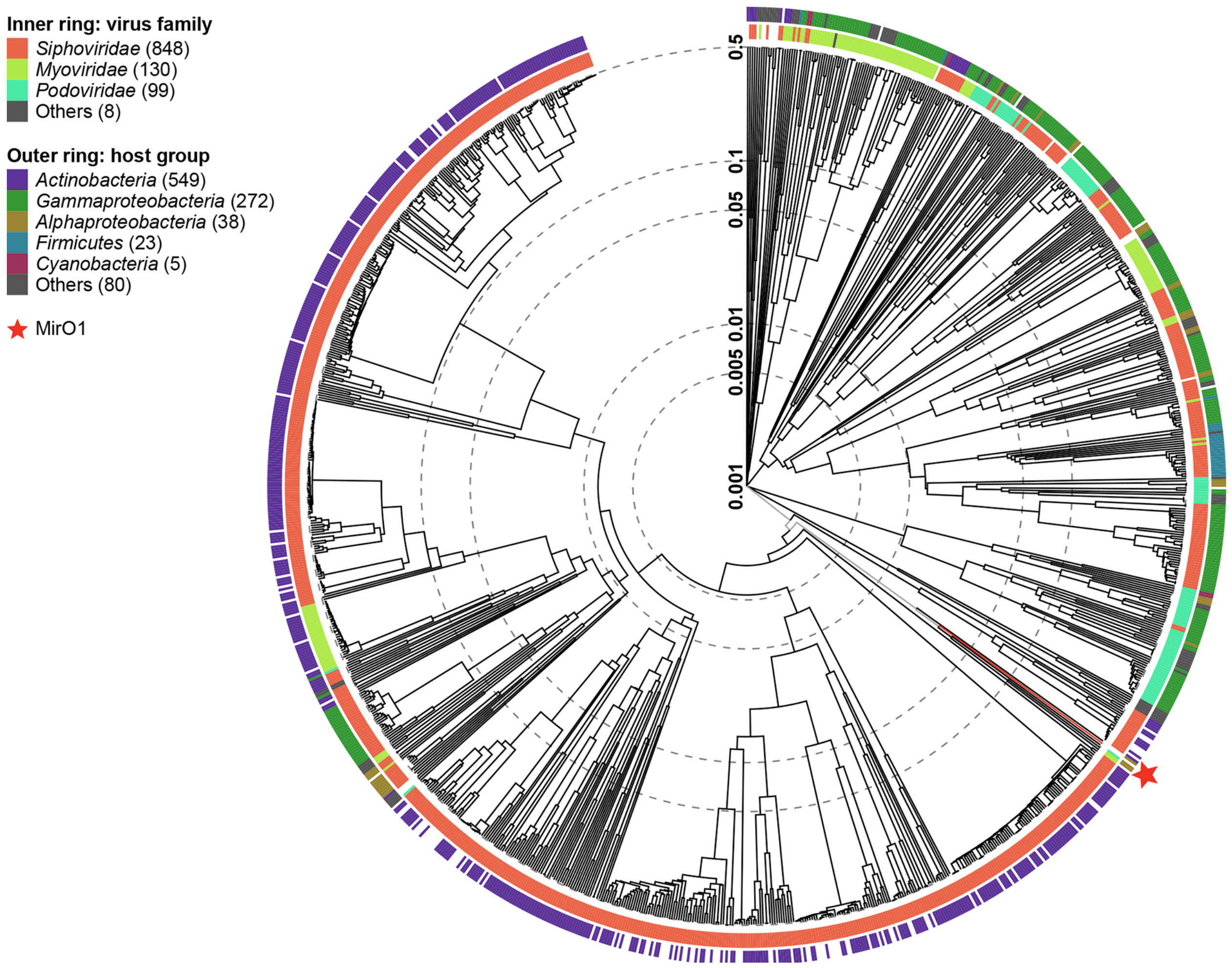


**Supplementary Figure S8.**

Phylogenetic analysis of the entire induced phage MirO1 genome.
